# RNF185 destabilizes specific membrane proteins that span all ERAD branches

**DOI:** 10.64898/2026.09.23.753758

**Authors:** Haruka Chino, Isabella M. Ruiz, Joshua H. Corbo, Joao A. Paulo, Sichen Shao

## Abstract

The endoplasmic reticulum (ER) is the primary site of eukaryotic membrane protein synthesis and quality control, which largely relies on ER-associated degradation (ERAD) to eliminate aberrant nascent proteins. In yeast, two ubiquitin ligases target proteins for ERAD based on aberrancies in substrate transmembrane (ERAD-M), lumenal (ERAD-L), or cytosolic (ERAD-C) domains. How an expanded repertoire of mammalian ERAD factors selects substrates across these classifications is unclear. Here, we show that the human ER-resident RNF185 ubiquitin ligase complex destabilizes a small but specific set of membrane proteins that span all three ERAD branches. Comparisons of three single-pass membrane proteins destabilized by RNF185 identify ERAD-M features in misoriented CHST10, ERAD-C features in unassembled SRPRB, and N-linked glycosylation-dependent ERAD-L features in ATP1B2. Our findings identify RNF185-destabilized membrane proteins with distinct aberrancies that collectively encompass all canonical ERAD substrate classifications and unexpectedly diverse quality control defects.

## Introduction

The endoplasmic reticulum (ER) is the major site for membrane protein synthesis, folding, which requires correct orientation of each transmembrane segment (TM) inserted into the lipid bilayer, posttranslational modifications, and complex assembly. Nascent proteins that fail biosynthesis are typically eliminated by ER-associated degradation (ERAD), which has three fundamental requirements^1,2^. First, substrate proteins must be recognized and ubiquitylated by ER-embedded ubiquitin ligase complexes, which may involve partial ‘retrotranslocation’ of the substrate across the ER membrane to access cytosolic ubiquitylation machinery. Second, ubiquitylated substrates are unfolded and extracted into the cytosol. This process requires the p97 AAA ATPase, its ubiquitin-binding adaptors, and is usually facilitated by membrane-embedded ERAD factors, including ubiquitin ligase complexes, that reduce the energetic barrier of protein extraction^1,3–5^. Finally, the substrates are subject to proteasomal degradation.

How ER ubiquitin ligase complexes select ERAD substrates is an outstanding question. Budding yeast, a major model system for studying ERAD, have two conserved RING-type ubiquitin ligases, Hrd1p (HRD1 in humans) and Doa10p (MARCH6 in humans), in the bulk ER. These ligases target proteins for ERAD based on functional ‘branches’ defined by the location of the substrate degron^6–8^. Hrd1p complexes handle ERAD-M and ERAD-L substrates that contain aberrant feature(s) in transmembrane or lumenal substrate domains, respectively. Although both ERAD-M and ERAD-L require Hrd1p, these two branches are distinguished by the requirement for additional cofactors to degrade ERAD-L but not ERAD-M substrates and separation-of-function Hrd1p mutants^6,9–11^. Doa10p handles ERAD-C substrates with aberrant features in cytosolic domains and select ERAD-M substrates, notably tail-anchored proteins with a single C-terminal TM that are otherwise cytosolic^6–8,12–14^. Membrane proteins can be substrates of any branch based on degron localization, and this framework for ERAD substrate selection has been proposed to be generalizable^6^. However, in contrast to yeast, over ten different ubiquitin ligases have been implicated in ERAD in human cells^1,2,15,16^. How these complexes select substrates in relation to the three canonical ERAD branches is unclear.

In this study, we systematically identify the determinants underlying the destabilization of three single-pass membrane proteins: a Golgi-resident carbohydrate sulfotransferase CHST10, the membrane-anchored signal recognition particle (SRP) receptor subunit SRPRB, and a sodium-potassium transporter subunit ATP1B2, by the metazoan ER-resident ubiquitin ligase RNF185. Although RNF185 and its obligate binding partner, membralin (MBRL, also known as TMEM259) are linked to ERAD-M^17–20^, we identify distinct substrate protein quality control defects that map onto all three branches of ERAD. Selective destabilization of CHST10 upon TM misorientation is consistent with ERAD-M, ‘orphan’ SRPRB that fails to assemble with its binding partner SRPRA displays ERAD-C features, and N-linked glycosylation requirements impart ERAD-L properties to ATP1B2. The revelation that RNF185 destabilizes specific proteins with degrons spanning membrane, cytosolic, and lumenal domains reshapes the conceptual framework for studying ERAD substrate selection.

## Results

### The RNF185 complex specifically destabilizes misoriented CHST10

CHST10 has a single N-terminal TM prone to membrane insertion in the wrong orientation^13^. Functional CHST10 has a ‘type II’ topology, which places its 6-residue N-terminal domain (NTD) in the cytosol and its catalytic C-terminal domain (CTD) in the ER, and ultimately the Golgi, lumen. CHST10 tends to misinsert owing to a lack of N-terminal positive charges, which are preferentially retained in the cytosol according to the ‘positive-inside’ rule^13,21,22^. We previously showed that the protein dislocation activity of the ER-resident P5A-ATPase ATP13A1 corrects CHST10 topology^13,23^. In ATP13A1 knockout (KO) cells, CHST10 is predominantly misoriented and degraded through incompletely defined ERAD mechanisms^13^ (Figure 1A).

**Figure 1.**
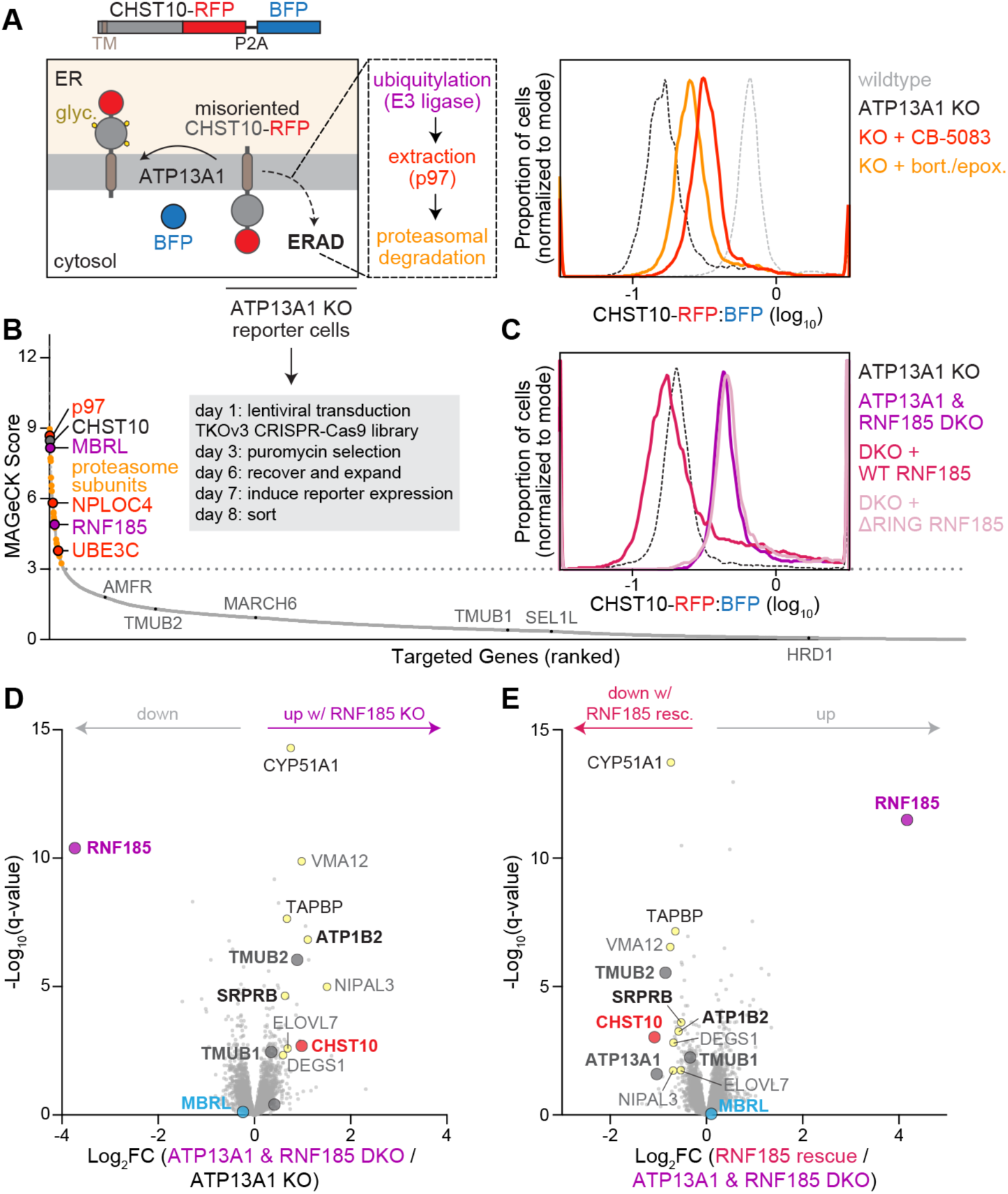
The RNF185 complex selectively destabilizes misoriented CHST10. **(A)** Misoriented CHST10 undergoes ER-associated degradation (ERAD). Scheme (left) and fluorescent flow cytometry histogram (right) of the RFP:BFP ratios of wildtype (gray) or ATP13A1 knockout (KO, black) Flp-In 293 T-REx cells expressing a stability reporter of CHST10 without or with 6 hr inhibition of p97 AAA-ATPase (1 µM CB-5083, dark orange) or proteasome (0.5 µM bortezomib, bort., and 0.5 µM epoxomicin, epox., light orange) activity. **(B)** A genome-wide CRISPR knockout screen on ATP13A1 KO cells expressing the CHST10 stability reporter identifies gene targets of sgRNAs enriched, by rank order, in cells with high RFP:BFP ratios. Proteasome subunits (light orange), CHST10, and other ubiquitin-proteasome system factors (dark orange or gray) are indicated. **(C)** Fluorescent flow cytometry of the CHST10 stability reporter in ATP13A1 and RNF185 DKO cells without (purple) or with re-expression of wildtype (WT) RNF185 (hot pink) or RNF185 lacking its catalytic RING domain (ΔRING, lavender). **(D)** RNF185 deletion stabilizes endogenous misoriented CHST10. TMT-MS volcano plot showing changes in protein levels upon knocking out RNF185 (DKO) in ATP13A1 KO cells. **(E)** TMT-MS volcano plot showing changes in protein levels upon re-expressing Flag-tagged RNF185 (rescue) in ATP13A1 and RNF185 DKO cells.

To study misoriented CHST10 ERAD, we established a dual fluorescent protein stability reporter system in Flp-In 293 T-REx cells comprising CHST10 with a C-terminal mCherry (referred to as RFP) tag, separated from TagBFP (referred to as BFP) by a P2A ribosome skipping sequence (Figure 1A). Because BFP is synthesized from the same transcript as CHST10-RFP, the RFP:BFP ratio is a readout for posttranslational CHST10 stability. As expected, the RFP:BFP ratio of this reporter is lower in ATP13A1 KO cells than in wildtype cells and is stabilized with p97 or proteasome inhibition, consistent with ERAD (Figure 1A).

To identify the factors that degrade misoriented CHST10, we performed a genome-wide screen using the CHST10 stability reporter in ATP13A1 KO cells and the Toronto KnockOut CRISPR-Cas9 library (TKOv3), which contains 70,948 sgRNAs targeting 18,053 genes, averaging 4 sgRNAs per gene (Figure 1B). Compared to untransduced ATP13A1 KO cells, transduced ATP13A1 KO cells displayed a population with higher RFP:BFP ratios, similar to the levels in wildtype cells (Figure S1A). This population was enriched with sgRNAs targeting factors required for ERAD, including proteasomal subunits, p97, and the p97 adaptor NPLOC4 (Figure 1B, Table S1). sgRNAs against CHST10 were also enriched, suggesting that CHST10-RFP levels increase to compensate for decreased endogenous CHST10 levels.

The top ubiquitin ligase hit was RNF185, a single-pass ER-resident RING-type ligase (Figure 1B). MBRL, a multi-spanning binding partner of RNF185 required for RNF185 stability^17^, ranked even higher (Figure 1B, S1B). Indeed, knocking out RNF185 or MBRL, but not another ER ubiquitin ligase AMFR, stabilized CHST10-RFP in ATP13A1 KO cells (Figure 1C, S1C). Importantly, re-expressing wildtype (WT) RNF185 but not a mutant lacking the catalytic RING domain (ΔRING) in ATP13A1 and RNF185 double knockout (DKO) cells decreased RFP:BFP ratios to levels comparable to ATP13A1 KO cells (Figure 1C). While a targeted knockdown approach previously implicated RNF185 and a partial role for AMFR in destabilizing misoriented CHST10^13^, the role of AMFR was not reproduced in our screen or with KO cells. Based on the results of unbiased genome-wide screening and the specific rescue of CHST10 destabilization by WT RNF185, we conclude that RNF185 is the primary ubiquitin ligase that mediates misoriented CHST10 ERAD.

Supporting this conclusion, unbiased proteomic profiling revealed that knocking out RNF185 or MBRL stabilized endogenous CHST10 in ATP13A1 KO but not wildtype cells, and that re-expressing RNF185 in ATP13A1 and RNF185 DKO cells rescued CHST10 destabilization (Figure 1D,E, S1D,E, Table S2, S3). The selective destabilization of CHST10 in ATP13A1 KO backgrounds is distinct among previously and newly-identified RNF185 substrates (discussed below) and supports the claim that RNF185 specifically eliminates misoriented CHST10 that would otherwise accumulate in ATP13A1 KO cells.

RNF185 and MBRL also function with TMUB1/2^17^: paralogous membrane-anchored proteins with a cytosolic ubiquitin-like (Ubl) domain that may recruit p97^24^ (Figure S1B). Both TMUB1 and TMUB2 are stabilized upon RNF185 deletion and destabilized upon re-expression of RNF185 (Figure 1D,E), consistent with proximity to degradation machinery. We hypothesized that the TMUB paralogs were not identified in our screen because they function redundantly. Indeed, while knocking down TMUB1 or TMUB2 individually did not change CHST10-RFP levels in ATP13A1 KO cells, simultaneously depleting both stabilized CHST10-RFP (Figure S1F,G). Thus, all factors implicated in the RNF185 ERAD pathway are required to destabilize CHST10 in the absence of ATP13A1.

### Misorientation is sufficient for CHST10 destabilization by the RNF185 complex

Re-expressing ATP13A1 in ATP13A1 KO cells rescues correctly oriented CHST10, which can be assayed by the modification of N-linked glycosylation sites in the CHST10 CTD that result in an increased molecular weight^13^. Because N-linked glycosylation machinery resides exclusively in the ER lumen, these modifications can only occur when CHST10 is correctly oriented. In contrast, knocking out RNF185 or MBRL in ATP13A1 KO cells predominantly stabilized unmodified, misoriented CHST10 (Figure 2A).

**Figure 2.**
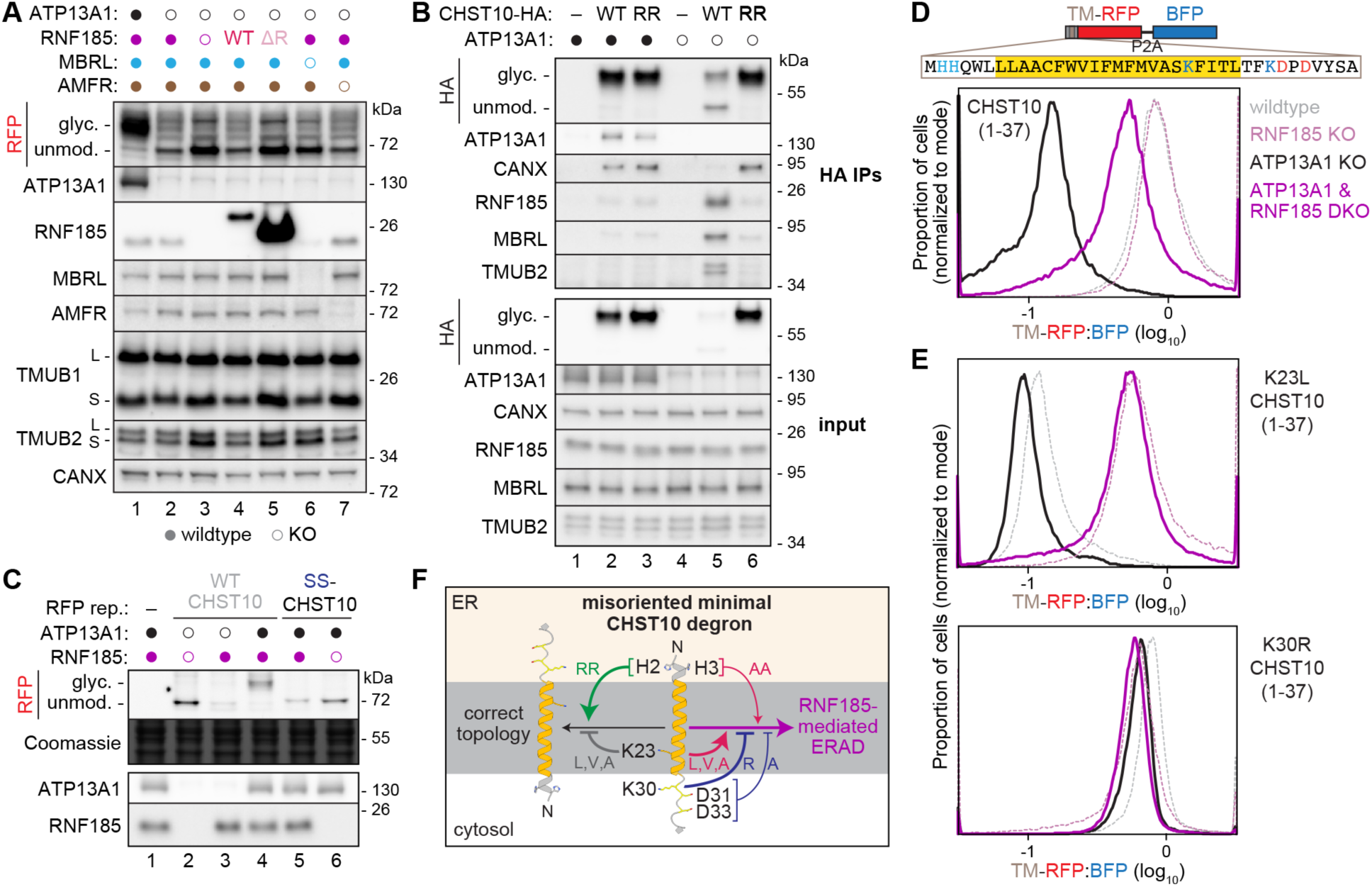
Misorientation of the CHST10 TM is sufficient for destabilization by RNF185. **(A)** SDS-PAGE and immunoblotting of lysates from cells without (filled circles) or with (open circles) knockout (KO) of the indicated factor and re-expression of wildtype (WT) or ΔRING (ΔR) RNF185. Glycosylated (glyc.) and unmodified (unmod.) CHST10, and long (l) and short (s) TMUB isoforms are indicated. **(B)** SDS-PAGE and immunoblotting of lysates before (bottom) or after anti-HA immunoprecipitations (IPs, top) of WT or H2R/H3R (RR) HA-tagged CHST10 expressed in wildtype (filled circles) or ATP13A1 KO (open circles) cells. **(C)** Forcing CHST10 misorientation by fusing the N-terminal cleavable signal sequence (SS) of preprolactin is sufficient for destabilization. SDS-PAGE and in-gel RFP fluorescence and Coomassie staining (top) or immunoblotting (bottom) of lysates from cells expressing the WT CHST10 or the SS-CHST10 stability reporter without (filled circles) or with ATP13A1 (black open circles) and/or RNF185 (purple open circles) KO. **(D)** The transmembrane (TM) region of CHST10 is sufficient for ERAD. Scheme (top) and fluorescent flow cytometry (bottom) of a stability reporter of the first 37 amino acids of CHST10 (TM-RFP) in wildtype, RNF185 KO, ATP13A1 KO, or ATP13A1 and RNF185 double knockout (DKO) Flp-In 293 T-REx cells. The 37 amino acid sequence shows the TM (yellow), basic (blue), and acidic residues (red). **(E)** As in (D) of the CHST10 TM reporter with K23L (top) or K30R (bottom) mutations. **(F)** Summary of mutations that promote (green arrows) or prevent (gray lines) correct CHST10 topology (black arrow), or that enhance (pink arrows) or inhibit (navy lines) CHST10 destabilization by RNF185 (purple arrow).

Misoriented CHST10 also specifically interacts with the RNF185 complex, as assayed by affinity purifications of C-terminally HA-tagged CHST10. In wildtype cells, correctly oriented CHST10-HA pulled down the ER protein chaperones calnexin and BiP (Figure 2B: lane 2, S2A: lane 1). In ATP13A1 KO cells, misoriented CHST10-HA specifically enriched for RNF185, MBRL, and TMUB2 (Figure 2B: lane 5, S2A: lanes 2-4). Mutating the histidines in the CHST10 NTD to arginines (H2R/H3R, referred to as RR-CHST10), which enforces correct CHST10 orientation in ATP13A1 KO cells^13^, eliminated RNF185 complex interaction and restored CHST10 glycosylation and ER chaperone interaction (Figure 2B: lanes 3 and 6, S2A: lane 5).

The interaction between misoriented CHST10, MBRL, and if present, a catalytically inactive C76A/C79A (CA) RNF185, was nearly stoichiometric in ATP13A1 and RNF185 DKO cells (Figure S2A: lanes 3 and 4). The persistence of the interaction between misoriented CHST10 and MBRL in the absence of RNF185 (Figure S2A: lane 3) is consistent with the model that MBRL bridges substrates and RNF185^18^. Misoriented CHST10 also pulled down near stoichiometric amounts of cytosolic HSP70 and HSP90 (Figure S2A: lanes 3 and 4), but inhibitors against these chaperones did not change CHST10 levels (Figure S2B). Thus, the RNF185 complex selectively engages and prevents the accumulation of misoriented CHST10.

We tested if misorientation is sufficient for CHST10 ERAD by fusing the cleavable signal sequence (SS) of the secreted protein preprolactin to the CHST10 stability reporter (referred to as SS-CHST10). Signal sequences engage the ER translocation in an N_cyto_-C_lum_ orientation^25^, which will initiate translocation of the CHST10 NTD into the ER lumen and force the TM to insert in the wrong orientation. Consistent with forced misorientation, SS-CHST10 was not glycosylated (Figure 2C). In addition, SS-CHST10 levels were lower than WT CHST10 in wildtype cells and were stabilized by knocking out RNF185 (Figure 2C: lanes 4-6, S2C). SS-CHST10 also specifically interacted with the RNF185 complex (Figure S2D). Thus, RNF185 destabilizes SS-CHST10 in wildtype cells, analogous to the destabilization of misoriented CHST10 in ATP13A1 KO cells. These data show that the RNF185 complex selectively engages misoriented CHST10 for ERAD.

### The TM region of CHST10 is a minimal degron

To date, all RNF185 substrates analyzed are categorized as ERAD-M according to one of two parameters. The first is whether a ‘minimal degron’ containing the substrate TM region is sufficient for RNF185-dependent destabilization, as observed for several cytochrome P450 family members such as CYP26A1 and CYP51A1^17,19^. The second is whether a specific mutation within a substrate TM abolishes RNF185-mediated ERAD, as was shown for unassembled, or ‘orphan’, tapasin (TAPBP)^18^. To test these parameters, we generated a stability reporter of the TM region of CHST10 containing only the first 37 amino acids (Figure 2D). Like full-length CHST10, the minimal CHST10-TM reporter was destabilized in ATP13A1 KO cells, and its levels were restored by p97 and proteasome inhibitors or by knocking out RNF185 or MBRL (Figure 2D, S2E). The sufficiency of the CHST10 TM region for RNF185-mediated degradation is consistent with ERAD-M.

Mutating a lysine (K428) in the TM of TAPBP impairs RNF185-mediated destabilization^18^. CHST10 also contains an intramembrane lysine (K23). Surprisingly, mutating K23 to hydrophobic amino acids (A, V, or L), but not to arginine, independently destabilized both the full-length CHST10 and CHST10-TM reporter in wildtype cells to levels similar to those in ATP13A1 KO cells, and knocking out RNF185 stabilized the K23L mutant in all cases (Figure 2E, S2F-I). We hypothesize that these K23 mutants target CHST10 to ERAD even in wildtype cells by dominantly driving misorientation, as more hydrophobic N-terminal TMs tend to insert in the N_lum_-C_cyto_ topology^22^. Consistent with this idea, these K23 mutants showed less glycosylation (Figure S2G,I). Thus, unlike TAPBP, the intramembrane lysine in CHST10 is not necessary for RNF185-mediated degradation, although the destabilization of misoriented CHST10 K23 mutants in wildtype cells suggests that this intramembrane lysine is an important signal for ATP13A1-mediated topology correction (Figure 2F).

In contrast, mutating charged residues immediately following the TM (K30, D31, and D33), individually or in combination, stabilized both the CHST10-TM reporter in ATP13A1 KO cells and the more topologically resistant SS-CHST10 reporter in wildtype cells (Figure 2E, S2H-K). A triple (K30A/D31A/D33A) and, unexpectedly, a single K30R SS-CHST10 mutant were most strongly stabilized and insensitive to RNF185 (Figure S2J,K). In comparison, mutating the two histidines in the NTD (H2A/H3A) further destabilized SS-CHST10 in an RNF185-dependent manner. K30A and K30R mutations also stabilized the CHST10-TM reporter (Figure 2E, S2H,I). Two non-exclusive possibilities may contribute to these K30 mutant effects. First, the higher pKa of arginine may better satisfy the positive-inside rule, implicating RNF185-mediated ERAD-M in positive-inside rule enforcement. Second, mutating K30 may eliminate an important ubiquitin acceptor site, although the partial effect of the K30A SS-CHST10 mutant argues against this as the sole explanation (Figure S2J). Collectively, these findings show that RNF185-mediated destabilization of CHST10 requires an N_lum_-C_cyto_ TM topology and is influenced by polar residues flanking the TM, particularly those near the cytosolic leaflet of the membrane (Figure 2F).

### The RNF185 complex specifically destabilizes orphan SRPRB

To further probe the substrate determinants of RNF185-mediated degradation, we analyzed two additional single-pass membrane proteins, SRPRB and ATP1B2, with uncharacterized ERAD pathways. Proteomics analysis revealed that RNF185 destabilizes these proteins in both wildtype and ATP13A1 KO cells (Figure 1D,E, S3A-D, Table S2, S3). Like CHST10, SRPRB and ATP1B2 each have a short NTD and a folded CTD flanking their TM. These shared architectural features facilitate comparisons of the molecular basis for their destabilization.

SRPRB is a subunit of the heterodimeric signal recognition particle (SRP) receptor, the membrane docking site for SRP-bound ribosomes synthesizing nascent proteins with ER targeting signals. SRPRB has a 36-residue NTD that translocates into the ER, followed by a TM and a cytosolic C-terminal GTPase domain that recruits SRPRA, a soluble GTPase^26^. This topology resembles misoriented CHST10 and RNF185-destabilized cytochrome P450 proteins^17,19^. Consistent with our proteomics results, immunoblotting showed that endogenous SRPRB levels increased upon RNF185 deletion and were restored by re-expression of WT but not ΔRING RNF185 (Figure 3A). A stability reporter of SRPRB was similarly regulated by RNF185 (Figure 3B, S3E,F). Thus, the ubiquitin ligase activity of RNF185 destabilizes SRPRB.

**Figure 3.**
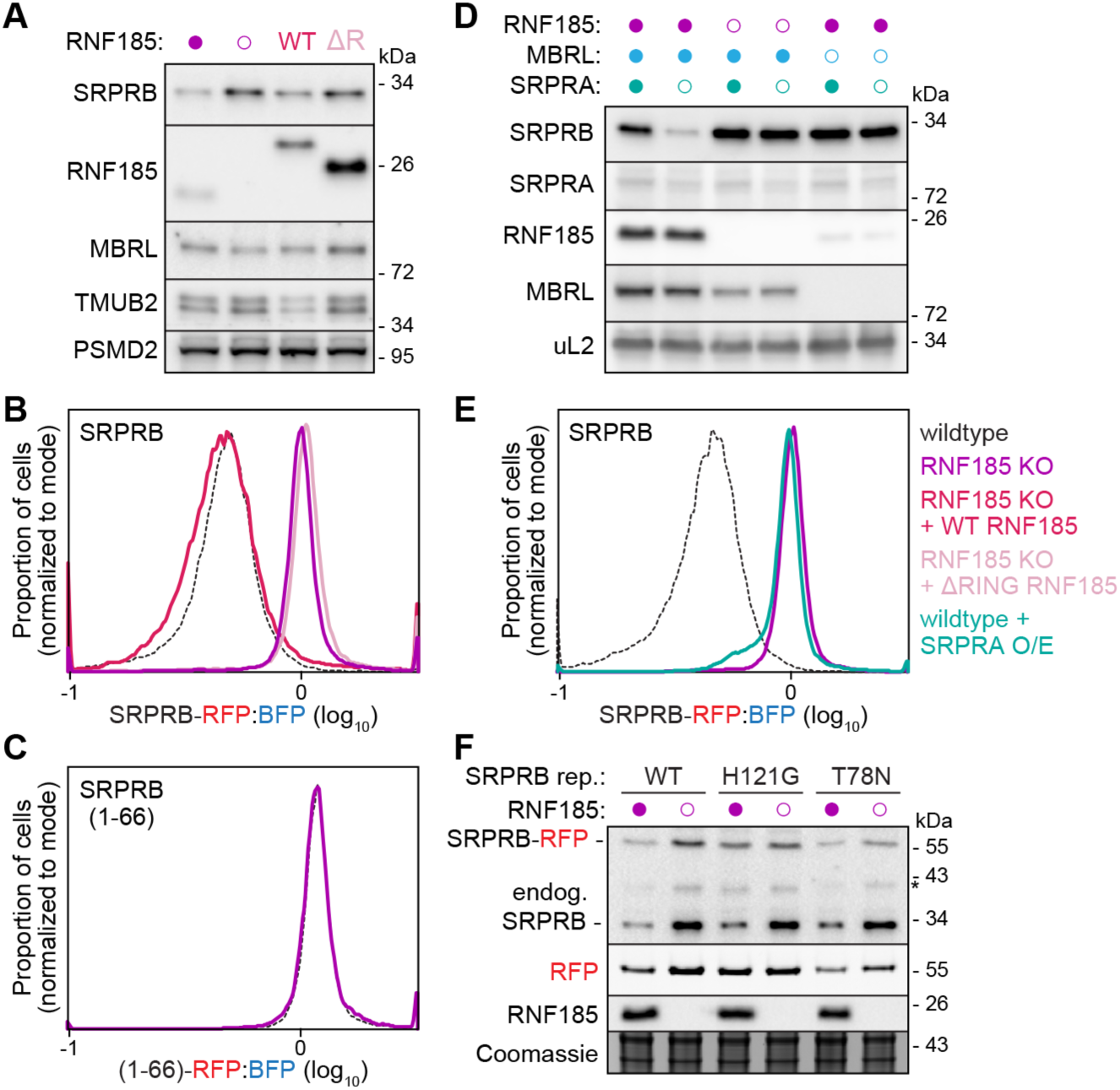
RNF185 selectively destabilizes orphaned SRPRB. **(A)** RNF185 deletion stabilizes endogenous SRPRB. Immunoblotting of lysates from wildtype (filled circle) or RNF185 knockout (KO) Flp-In 293 T-REx cells without (open circle) or with re-expression of wildtype (WT) or ΔRING (ΔR) Flag-tagged RNF185. **(B)** RFP-tagged SRPRB is stabilized in RNF185 KO cells (purple) compared to wildtype cells (black), assayed by fluorescent flow cytometry. Re-expression of WT (hot pink) but not ΔRING RNF185 (lavender) in RNF185 KO cells destabilizes SRPRB. **(C)** RNF185 KO (purple) does not change the levels of an RFP-tagged stability reporter of the first 66 amino acids of SRPRB, which includes the N-terminal transmembrane region (TM), assayed by fluorescent flow cytometry. **(D)** RNF185 destabilizes SRPRB orphaned from SRPRA. Immunoblotting as in (A) of cells without (filled circles) or with KO of RNF185 (purple open circles) or MBRL (blue open circles), or knockdown (KD) of SRPRA (open teal circles). **(E)** SRPRA overexpression stabilizes SRPRB. Fluorescent flow cytometry of the SRPRB stability reporter without or with overexpression (O/E) of SRPRA (teal). **(F)** Immunoblotting of wildtype (closed purple circles) or RNF185 KO (open purple circles) cells expressing RFP-tagged SRPRB stability reporters without (WT) or with GTP-locked (H121G) or GDP-locked (T78N) mutations.

Unexpectedly, unlike CHST10 (Figure 2D) and RNF185 substrates in the cytochrome P450 protein family^17,19^, a minimal stability reporter containing only the TM region of SRPRB (residues 1-66) was insensitive to RNF185 (Figure 3C, S3G). This raised the possibility that the cytosolic CTD may be important for RNF185-mediated destabilization. Because the SRPRB CTD binds SRPRA, we queried how depleting SRPRA impacts SRPRB. Knocking down SRPRA reduced SRPRB levels (Figure 3D: lanes 1 and 2), revealing specific destabilization of ‘orphan’ SRPRB. Importantly, SRPRA depletion did not destabilize SRPRB in RNF185 or MBRL KO cells (Figure 3D: lanes 3-6), suggesting that the RNF185 complex specifically degrades orphan SRPRB.

Further supporting a role for RNF185 in SRPRB orphan protein quality control, overexpressing SRPRA stabilized the SRPRB stability reporter to levels comparable to RNF185 KO (Figure 3E, S3H). In addition, consistent with reports that the SRPRB-SRPRA interaction is favored when both are bound to GTP^26,27^, the levels of a GTP-locked (H121G) SRPRB mutant were higher than WT SRPRB and less sensitive to RNF185 deletion (Figure 3F, S3I). In contrast, endogenous SRPRB in the same cells and a GDP-locked (T78N) SRPRB mutant remained sensitive to RNF185 (Figure 3F). These data indicate that RNF185 specifically destabilizes orphan SRPRB based on ERAD-C features unmasked in the absence of its cytosolic binding partner.

### Lumenal N-linked glycosylation sites destabilize ATP1B2 via RNF185

We next investigated ATP1B2, a beta subunit of sodium-potassium transporters (Figure S4A). Like CHST10, ATP1B2 is a type II protein: it has a 39-residue cytosolic NTD followed by a TM and a globular CTD that should translocate into the ER and ultimately reside on the extracellular side of the plasma membrane. Validating RNF185-mediated destabilization, an RFP-tagged ATP1B2 reporter was stabilized by ERAD inhibitors and in RNF185 KO cells, where it was destabilized upon re-expression of WT but not ΔRING RNF185 (Figure 4A,B, S4B).

**Figure 4.**
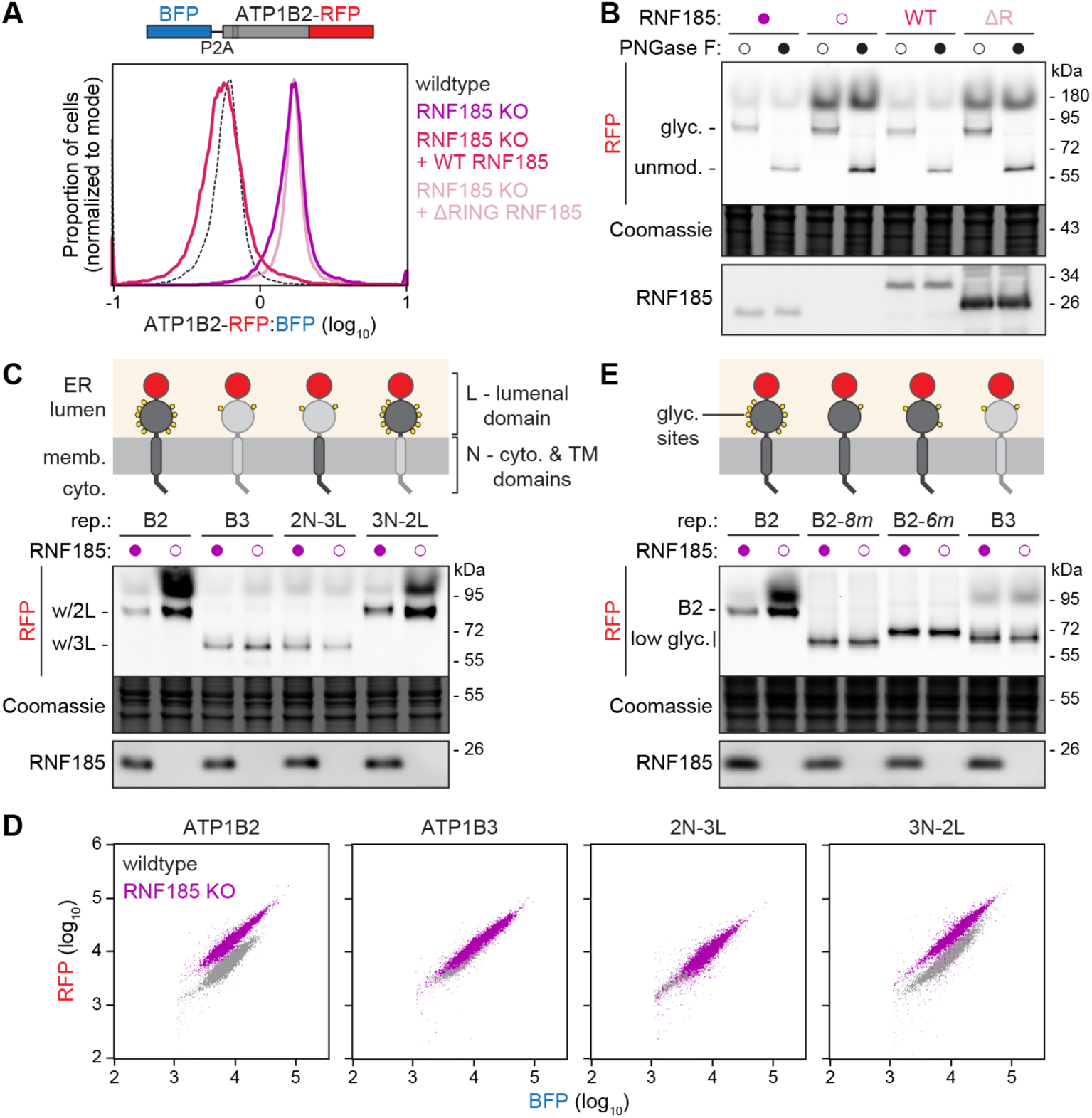
RNF185 selectively destabilizes N-linked glycosylated ATP1B2. **(A)** Fluorescent flow cytometry of wildtype (black) or RNF185 knockout (KO) Flp-In 293 T-REx cells expressing a stability reporter of RFP-tagged ATP1B2 without (purple) or with re-expression of wildtype (WT, hot pink) or ΔRING (lavender) RNF185. **(B)** Correctly oriented ATP1B2 is stabilized in RNF185 KO cells. SDS-PAGE and in-gel RFP fluorescence and Coomassie staining (top) or immunoblotting (bottom) of cells as in (A) before or after treatment with PNGase F to remove N-linked glycans. **(C)** The ATP1B2 lumenal domain confers sensitivity to RNF185. SDS-PAGE analysis as in (B) of lysates from wildtype (filled circles) or RNF185 KO (open circles) cells expressing RFP-tagged stability reporters of ATP1B2, ATP1B3, or chimeras of the N-terminal cytosolic and TM domain and C-terminal lumenal domain of the two paralogs as indicated. **(D)** Fluorescent flow cytometry scatter plots showing RFP vs. BFP levels of the ATP1B2, ATP1B3, or the chimeric reporters as in (C). **(E)** Mutating N-linked glycosylation sites in ATP1B2 abolishes sensitivity to RNF185. SDS-PAGE analysis as in (C) of ATP1B2 reporters with mutations to disrupt 8 or 6 N-linked glycosylation motifs.

A commonality of all single-pass RNF185 substrates analyzed so far is a N_lum_-C_cyto_ TM topology^17–19,28^ (Figure 1-3). Considering our findings with CHST10, we asked whether RNF185 destabilizes misoriented ATP1B2. However, treatment with the glycosidase PNGase F revealed that RNF185 predominantly destabilizes glycosylated ATP1B2 (Figure 4B). Because all possible N-linked glycosylation motifs of ATP1B2 reside in its CTD and can only be modified in the ER lumen, this finding and the observation that RNF185 deletion does not selectively stabilize unmodified ATP1B2 suggest that RNF185 destabilizes correctly oriented, type II ATP1B2. An N_cyto_-C_lum_ TM topology is a distinction among RNF185 substrates that have been studied.

Like SRPRB, a minimal stability reporter containing only the ATP1B2 NTD and TM (residues 1-79) was insensitive to RNF185 (Figure S4C), raising the intriguing possibility that the lumenal CTD of ATP1B2 is important for destabilization. To test this possibility, we leveraged the similarity between ATP1B2 and its paralogs (Figure S4A) to identify the distinctive features of ATP1B2 required for destabilization. ATP1B1 and ATP1B3 were both insensitive to RNF185 and MBRL in our proteomics datasets (Table S2, S3). RNF185 KO also did not stabilize a reporter of ATP1B3, which is 48% identical to ATP1B2 (Figure 4C,D). Chimeras comprising the NTD and TM of one paralog attached to the CTD of the other revealed that RNF185 specifically destabilized reporters containing the ATP1B2 CTD. Thus, consistent with ERAD-L, the lumenal CTD of ATP1B2 contains the critical information for RNF185-mediated destabilization.

Although the ATP1B2-ATP1B3 chimeric reporters differ by no more than 11 amino acids in length, those containing the ATP1B2 CTD were not only specifically destabilized by RNF185 but also migrated ∼15 kDa higher by SDS-PAGE (Figure 4C, S4A). This difference reflects the presence of nine potential NxS/T glycosylation motifs in the ATP1B2 CTD versus two in the ATP1B3 CTD. To test if N-linked glycosylation impacts protein stability, we mutated six (ATP1B2-*6m*) or eight (ATP1B2-*8m*) NxS/T motifs in ATP1B2 (Figure S4A). Both ATP1B2 loss-of-glycosylation mutants migrated at lower molecular weights, were more stable than WT ATP1B2 in wildtype cells, and were not further stabilized by RNF185 deletion (Figure 4E, S4D), although adding five glycosylation sites to ATP1B3 (ATP1B3+*5g*) was not sufficient to confer RNF185 sensitivity (Figure S4E,F). As ATP1B1 and ATP1B3 are both at least an order-of-magnitude more abundant than ATP1B2 in dividing cells^29,30^, prolonged glycosylation-coupled protein folding in the ER lumen may provide a specific means for ATP1B2 abundance regulation by the RNF185 complex. The importance of lumenal N-linked glycosylation for ATP1B2 degradation, considered together with our findings with CHST10 and SRPRB, establishes that the RNF185 complex destabilizes membrane proteins that span all three ERAD branches.

## Discussion

For two decades, ERAD-M, -L, and -C have proved useful for classifying ERAD substrates. Yet our dissection of RNF185-destabilizing features suggests that these categories do not generally define the substrate specificities of mammalian ERAD ligases. The concept of functional ERAD ‘branches’ emerged from the distinct substrate specificities of Hrd1p and Doa10p in yeast^6–8^, and it seems reasonable that substrate selection by ERAD ligases would segregate based on degron location. Several ER-resident ligases are reported to destabilize proteins representing two of the three canonical ERAD branches, with at least one being ERAD-M^6–14,31,32^. However, RNF185 specifically destabilizes proteins with quality control defects that span lumenal, membrane, and cytosolic domains. To our knowledge, RNF185 is the first ER ubiquitin ligase shown to act across all three ERAD branches based on two rigorous benchmarks: first, evidence that RNF185 destabilizes endogenous protein levels, and second, identification and, importantly, experimental validation of specific substrate mutations that abolish RNF185-mediated destabilization.

Based on these criteria, we show that three membrane proteins, each with a single N-terminal TM and a globular CTD, are destabilized by RNF185 and encompass all three ERAD classifications. In addition, these proteins present distinct issues involving protein topology (misoriented CHST10), complex assembly (orphan SRPRB), and posttranslational modification (glycosylated ATP1B2) that trigger their destabilization by RNF185. Thus, RNF185 is also not dedicated to a specific protein quality control purpose. Other proteins destabilized by RNF185 include multi-spanning membrane proteins^33–35^ and peripheral proteins^36^ (Figure S3C). The molecular determinants underlying their destabilization remain to be determined.

RNF185 is probably not the only ERAD ligase that destabilizes proteins with diverse degrons, and additional RNF185 substrates may emerge under specific conditions, as exemplified by CHST10, which is destabilized only in the absence of ATP13A1 (Figure 1). RNF185 simultaneously displays remarkable selectivity, as it does not destabilize many proteins that closely resemble its substrates, including the paralogs of ATP1B2 (Figure 4C,D) and most cytochrome P450 family members^17,19^. Moreover, RNF5, a paralog of RNF185 that may act redundantly in some contexts^33,34^, does not effectively compensate for RNF185 loss for the substrates examined here. While the basis of this specificity is unclear, RNF5, RNF185, and MBRL have all been implicated in viral responses^28,37–39^, raising the possibility that their substrate preferences emerged from or co-evolved with pathogenic threats.

An essential protein for neural homeostasis, MBRL interacts with and stabilizes RNF185 but not RNF5^17,20,40^. MBRL may bridge RNF185 and at least some ERAD-M substrates because MBRL-substrate interactions persist when RNF185 is absent, and MBRL overexpression impairs substrate degradation^17,18^, consistent with a separation of MBRL-substrate and MBRL-RNF185 complexes. However, the molecular basis of MBRL-substrate interaction remains to be determined. It is thus unknown if MBRL is sufficient to select RNF185 substrates, particularly those destabilized based on ERAD-C or ERAD-L features. Interestingly, MBRL overexpression rescued SRPRB destabilization in conditions that failed to rescue CHST10 and ATP1B2 destabilization (Figure S1E, S3D). This indicates that MBRL may be required only to stabilize RNF185, and not for SRPRB binding, for SRPRB degradation, although it is not yet clear if other factors play a role in substrate selection. Identifying such auxiliary requirements awaits future functional genetic analysis and biochemical reconstitutions.

## Acknowledgements

We thank G. Nelson for help with bioinformatics, J. Zhang for help with cell sorting, and M. McKenna, J. Coelho, D. Sherpa, M. Rale, and other Shao Lab members for useful discussions. Sequencing was performed at the Harvard Bauer Core Facility. This work was supported by a Takeda Science Foundation fellowship (HC), a National Science Foundation graduate research fellowship (JHC), National Institutes of Health R35GM156406 (JAP), National Institutes of Health R01AG073277 (SS), and a Smith Family Foundation Odyssey Award (SS). SS is an Investigator with the Howard Hughes Medical Institute.

## Author contributions

Conceptualization: HC, IR, SS; Investigation: HC, IR, JHC, JAP, SS; Funding acquisition: JAP, SS; Supervision: JAP, SS; Writing – original draft: HC, SS; Writing – review & editing: HC, IR, JHC, JAP, SS

## Declaration of interests

The authors declare no competing interests.

## Data availability

Sequencing data were deposited in the NCBI Gene Expression Omnibus (GEO) with accession code GSE338468. Proteomics data were deposited in the Proteomics Identifications Database (PRIDE) with accession codes PXD080176 and PXD080177.

## Materials and Methods

### Plasmids and antibodies

Guide RNA sequences were inserted into pX458, lentiCRISPRv2 Puro or lentiCRISPRv2 Neo using standard molecular biology techniques. Stability and HA-tagged reporters, as well as FLAG-RNF185 and MBRL-FLAG, were assembled in pcDNA5/FRT/TO using standard molecular biology techniques and cDNA sequences from previously described CHST10 expression plasmids^13^ or pENTR-SRPRB, pENTR223-ATP1B2, pENTR223-ATP1B3, pDONR-RNF185, pDONR-MBRL (gifts from the Harper lab). For lentiviral production, FLAG-SRPRA, MBRL-FLAG, and FLAG-RNF185 were transferred into the pHAGE vector from the vectors described above or from pENTR-SRPRA (a gift from the Harper lab) by Gibson Assembly. Mutations were introduced using standard molecular biology techniques.

HRP-conjugated anti-FLAG M2 (Sigma A8592, 1:5000 for IB), HRP-conjugated anti-HA (Cell Signaling Technology 2999, 1:1000 for IB), anti-ATP13A1 (Proteintech 16244-1-AP, 1:2500 for IB), anti-CANX (Enzo Life Sciences ADI-SPA-865, 1:5000 for IB), anti-AMFR (Thermo Fisher 16675-1-AP, 1:5000 for IB), anti-RNF185 (Abcam ab181999, 1:5000 for IB), anti-TMUB1 (Abcam ab180586, 1:5000 for IB), anti-TMUB2 (Proteintech 28044-1-AP, 1:5000 for IB), anti-MBRL (Sigma-Aldrich HPA042669, 1:2500 for IB), anti-SRPRB (Proteintech 14636-1-AP, 1:5000 for IB), anti-uL2 (RPL8) (Abcam ab169538), and anti-PSMD2 (Bethyl A303-853A, 1:5000 for IB) antibodies were purchased. HRP-conjugated goat anti-rabbit (Jackson ImmunoResearch 111-035-003, 1:5000 for IB) secondary antibodies were also purchased. The anti-RFP (1:5000 for IB) and anti-SRPRA (1:2000 for IB) antibodies were gifts from the Hegde lab.

### Cell line generation, culture, and treatments

Parent HEK293T (ATCC: CRL-3216) and Flp-In 293 T-REx (Invitrogen: R78007) cell lines were maintained in DMEM with high glucose, GlutaMAX, and sodium pyruvate supplemented with 10% fetal bovine serum at 37°C and 5% CO_2_. To generate knockout cell lines, Flp-In 293 T-REx cells were transfected with pX458 with the target gRNA sequence using TransIT 293 (Mirus 2706) according to the manufacturer’s instructions. Two days after transfection, single GFP-positive cells were isolated by fluorescence-activated cell sorting (FACS) on a SONY SH800. Clonal lines were validated by immunoblotting and genotyping.

Cells constitutively expressing FLAG-SRPRA, MBRL-FLAG, or wildtype, ΔRING (Δ39-80), or CA (C76A/C79A) FLAG-RNF185 were generated using third generation lentiviral transduction. To generate lentiviral particles, HEK293T cells were transfected with a 2:1:1:1:10 ratio of pHDM-VSVG, pHDM-HGPM2, pHDM-tat1B, pRC-CMV-rev1B and pHAGE containing the desired insert using TransIT according to the manufacturer’s instructions. Cells constitutively expressing lentiCRISPRv2 Puro or lentiCRISPRv2 Neo gRNA were generated using second generation lentiviral transduction. To generate lentiviral particles, HEK293T cells were transfected with a 1:1:1 molar ratio of pMD2.G, psPAX2 and lenti-CRISPR v2 containing the desired gRNA using TransIT according to the manufacturer’s instructions. In all cases, the media was harvested 48 hr after transfection and used to transduce Flp-In 293 T-REx cells in the presence of 8 µg/mL polybrene (MOI 0.5). 48 hr after transduction, cells were placed under selection with 2 µg/mL puromycin or 1 mg/ml G418 for 48 hr. Validation was performed by immunoblotting for the FLAG-tagged proteins or knockdown targets.

All stability and HA-tagged reporters, as well as inducible rescue constructs, were integrated into the doxycycline-inducible Flp-In locus of the indicated Flp-In 293 T-REx cell lines by cotransfecting a 1:1 ratio of pOG44 and pcDNA5/FRT/TO containing the gene of interest using TransIT 293 according to the manufacturer’s instructions. Cells were selected with 2.5 µg/mL blasticidin and 50 µg/mL hygromycin for 2-3 weeks, and expression was validated by induction with 100 ng/mL doxycycline for 24 hr followed by immunoblotting or fluorescent flow cytometry.

Unless otherwise noted, reporter or rescue protein expression was induced with 100 ng/mL doxycycline for 24-48 hr. Cells were treated with 0.5 µM bortezomib (LC laboratories B-1408) and 0.5 µM epoxomicin (ApexBio 134381-21-8) for 6 hr to inhibit proteasomal activity, or 1 µM CB-5083 (Selleck S8101) for 6 hr to inhibit p97 activity. Cells were treated with 30 µM VER155008 (Millipore Sigma SML0271), 2 µM geldanamycin (Selleck S2713), or 500 nM SNX-0723 (MedChem express HY-119046) for 7 hr to inhibit HSP70 or HSP90. For siRNA-mediated knockdowns, reverse transfections were performed. Briefly, trypsinized cells were plated at 50% confluence into 6-well plates containing 30 pmol siRNA and Lipofectamine RNAiMAX according to manufacturer’s instructions 72 hr before analysis.

### Fluorescent flow cytometry

Flp-In 293 T-REx cells were harvested either directly in PBS or detached with trypsin-EDTA and collected in complete growth medium or 1% FBS/PBS. Cells pelleted by centrifugation at 500xg for 3 min were washed and resuspended in PBS or 1% FBS/PBS and filtered through a 35 µm mesh strainer prior to analysis. Data were collected on an Attune NxT flow cytometer and analyzed using FlowJo. Samples were gated for single cells, and fluorescence was excited with lasers at 405 nm (for TagBFP) and 561 nm (for mCherry3V), and collected using 440/50 and 615/20 emission filters, respectively. All flow cytometry experiments are representative of at least 3 independent replicates.

### CRISPR screen

A 1:1:1 molar ratio of the TKOv3 CRISPR/Cas9 library (Addgene 90294) and 2nd generation lentiviral packaging plasmids (pMD2.G and psPAX2) was transfected into HEK293T cells using TransIT according to the manufacturer’s instructions. After 18 hr, media containing transfection reagents was replaced by media with ViralBoost (ALSTEM VB100). After 72 hr, lentiviral particles were harvested and used to transduce 75 million ATP13A1-KO Flp-In 293 T-REx cells expressing the CHST10-mCherry-P2A-BFP stability reporter at an MOI of 0.3. Cells were grown for 48 hr and then selected with 2 μg/mL puromycin for 72 hr. 24 hr prior to sorting, 100 ng/mL doxycycline was added to induce expression of the stability reporter. 100 million unsorted cells were obtained for reference, and another 100 million cells were sorted for RFP^high^/BFP ratios comparable to wildtype Flp-In 293 T-REx cells expressing the CHST10 stability reporter (10%) using a SONY SH800. Samples were gated for single cells, and fluorescence was excited with lasers at 405 nm (for TagBFP) and 561 nm (for mCherry3V), and collected using 450/50, and 600/60 emission filters, respectively.

Genomic DNA was extracted using a BloodMaxi kit (QIAGEN 51194) for reference samples or a BloodMini kit (QIAGEN 51106) for sorted samples, according to the manufacturer’s protocols. The gRNA regions were PCR amplified using NEBNext Ultra II Q5 Master Mix (NEB M0544) and the LCV2-F and LCV2-R primers^41^. The PCR reactions were pooled for a second PCR reaction to attach the indices and sequencing adapters using the i5 (D501, D502, D503, D504) and i7 (D701, D702, D704, D705) primers. The 200 bp band from the PCR reaction was excised and purified from a 2% TBE-agarose gel for sequencing on an Illumina NextSeq 1000. Gene rankings were obtained using MAGeCK 0.5.9.4^42^.

### Cell lysis and affinity purifications

For SDS-PAGE and in-gel fluorescence and immunoblotting analysis of total cell lysates, cells were either harvested in cold PBS or washed in cold PBS on the plate. Cell lysis was performed either after centrifugation at 500xg for 5 min at 4°C or on plate with 1% Triton X-100 or 1% CHAPS in RNC buffer [50 mM HEPES, pH 7.4, 100 mM KOAc, 5 mM Mg(OAc)_2_] with 1x complete EDTA-free protease inhibitor (PIC) and 1 mM DTT. After centrifugation at 12,000-21,300xg for 10 min at 4°C, the clarified lysates were normalized based on total protein concentration measured by absorbance at 280 nm or by Bradford assay. To detect glycosylation, denatured cell lysates were incubated with PNGase F (NEB P0704) at 37°C for 1 hr according to the manufacturer’s instructions. 2-20 µg lysates were analyzed by SDS-PAGE and in-gel fluorescence or immunoblotting.

For HA-tagged CHST10 affinity purifications, Flp-In 293 T-REx cells induced to express CHST10-HA with 100 ng/mL doxycycline for 24 hr in a 15-cm plate were harvested in cold PBS and centrifuged at 1000xg for 3 min at 4°C. Unless otherwise indicated, cells were crosslinked by resuspension into 1 mL PBS containing 0.1% paraformaldehyde (PFA) and rotated for 10 min at room temperature. Crosslinking was stopped with 100 μL 2.5 M glycine, 25 mM Tris pH 7.4 followed by centrifugation at 1000xg for 3 min at 4°C. The cell pellet was resuspended and incubated in 500 μL 0.02% digitonin in RNC buffer with 1x PIC and 1 mM DTT on ice for 10 min before centrifugation at 1000xg for 5 min at 4°C. The pellet containing organellar membranes was washed once in RNC buffer and solubilized in 500 μL 1% Triton X-100 in RNC buffer with 1x PIC and 1 mM DTT for 10 min on ice and then clarified by centrifugation at 12,000xg for 10 min at 4°C. The supernatants normalized to a total protein concentration of 5 µg/µL were subjected to immunoprecipitation with 10 μL packed anti-HA Magnetic beads (Thermo Fisher Scientific 88836) for 1 hr with rotation at 4°C, followed by three washes with 1 mL 1% Triton X-100 in RNC buffer with 1 mM DTT and elution with protein sample buffer.

### Multiplexed proteomics

Cells were washed twice with 1x PBS, harvested on ice using a cell scraper in 1x PBS, pelleted via centrifugation at 1,000xg for 5 min at 4°C, and washed with 1x PBS before resuspension in 8 M urea (Sigma U5128), 50 mM NaCl, 50 mM EPPS (Sigma E9502) and 1x PIC. After 10 s of sonication, lysed cells were pelleted, and the protein concentration of the clarified sample was determined using a BCA kit (Thermo Fisher Scientific 23225). Then, 100 µg of each sample was incubated for 30 min at 37°C with 5 mM TCEP (Thermo Fisher Scientific 77720) for disulfide bond reduction with subsequent alkylation with 20 mM iodoacetamide (Sigma I6125) for 20 min at room temperature followed by quenching with 15 mM DTT for 15 min under gentle shaking. MeOH-chloroform precipitation of samples was performed as follows: to each sample, four parts MeOH was added followed by vortexing, one part chloroform was added followed by vortexing, and finally three parts H_2_O was added. After vortexing, the suspension was centrifuged for 5 min at 14,000xg and the aqueous phase around the protein precipitate was removed using a loading tip. The precipitate was washed twice with MeOH and resuspended in 200 mM EPPS (pH 8.0) and digested with Trypsin/Lys-C (Promega V5073, 1:100) at 37°C overnight with gentle shaking.

150 µL of digested samples were labeled by adding 10 µL of TMT reagent (Thermo Fisher Scientific A52045; stock: 20 mg/ml in acetonitrile (ACN), Millipore Sigma 34851) together with 50 µL of ACN to yield a final ACN concentration of approximately 25% (v/v) for 1 hr at room temperature before quenching the reaction with hydroxylamine (Thermo Fisher Scientific 90115) at a final concentration of 0.2% (v/v). The TMTpro-labeled samples were pooled at a 1:1 ratio, resulting in a consistent peptide amount across all channels. Pooled samples were vacuum centrifuged for 1 hr at room temperature to remove ACN, followed by reconstitution in 1% formic acid (FA, Sigma 543804), desalting using C18 solid-phase extraction (200 mg, Sep-Pak, Waters WAT036820) and vacuum centrifugation until near dryness. We fractionated the pooled, labeled peptide sample using basic pH reversed-phase HPLC and an Agilent 1260 pump equipped with a degasser and a UV detector (wavelength set at 220 and 280 nm). Peptides were subjected to a 50-min linear gradient from 5% to 35% ACN in 10 mM ammonium bicarbonate pH 8 at a flow rate of 0.6 mL/min over an Agilent 300Extend C18 column (3.5 μm particles, 4.6 mm internal diameter and 220 mm in length). The peptide mixture was fractionated into 96 fractions, which were consolidated into 24 super-fractions, of which 12 nonadjacent fractions were analyzed. Samples were subsequently acidified with 1% FA and vacuum centrifuged to near dryness. Each super-fraction was desalted via StageTip, dried again via vacuum centrifugation and reconstituted in 10 µL 5% ACN, 5% FA for LC–MS/MS processing.

Mass spectrometric data were collected on an Orbitrap Ascend instrument coupled to a Vanquish Neo UHPLC. The 100-µm capillary column was packed with 35 cm of Accucore 150 resin (2.6 μm, 150 Å; Thermo Fisher Scientific) at a flow rate of 340 nL/min. Data were acquired at ∼90 min per fraction. For MS2-based quantification, the scan sequence began with an MS1 spectrum (Orbitrap analysis, resolution 60,000, mass range 350–1,350 Th, automatic gain control target 100%, maximum injection time 50 ms). The hrMS2 stage consisted of fragmentation by higher energy collisional dissociation (normalized collision energy 36%) and analysis using the Orbitrap (automatic gain control 200%, maximum injection time 100 ms, isolation window 0.6 Th, resolution 45,000). Data were acquired using the FAIMSpro interface with the dispersion voltage set to 5,000 V, the compensation voltages set at −40, −60 and −80 V, and the TopSpeed parameter set at 1 s per compensation voltage. For real-time search (RTS) MS3-based quantification, the scan sequence began with an MS1 spectrum (Orbitrap analysis, resolution 60,000, mass range 350–1,350 Th, automatic gain control target 100%, maximum injection time 50 ms). The MS2 stage consisted of fragmentation by collision-induced dissociation (normalized collision energy 35%) and analysis using the ion trap (automatic gain control 100%, maximum injection time 60 ms, isolation window 0.7 Th, turbo ion trap scan rate). The database was set to human and we limited the number of peptides per protein per fraction to 2 and to 10 synchronous precursor selection (SPS) ions. The MS3 stage consisted of fragmentation by higher energy collisional dissociation (normalized collision energy 55%) and analysis using the Orbitrap (automatic gain control 200%, maximum injection time 250 ms, isolation window 1.2 Th, resolution 45,000). Data were acquired using the FAIMSpro interface with the dispersion voltage set to 5,000 V, the compensation voltages set at −40, −60 and −80 V, and the TopSpeed parameter set at 1 s per compensation voltage.

Spectra were converted to mzXML via MSconvert. Database searching included all entries from the human UniProt reference database (downloaded August 2021). The database was concatenated with one composed of all protein sequences for that database in the reversed order. Searches were performed using a 50-ppm precursor ion tolerance for total protein-level profiling. For MS2-based quantification, the product ion tolerance was set to 0.03 Da, while for MS3-based quantification this value was set to 1 Da. TMTpro labels on lysine residues and peptide N termini (+304.207 Da), as well as carbamidomethylation of cysteine residues (+57.021 Da) were set as static modifications, while oxidation of methionine residues (+15.995 Da) was set as a variable modification. Peptide–spectrum matches (PSMs) were adjusted to a 1% false discovery rate. PSM filtering was performed using a linear discriminant analysis, as described previously^43^ and then assembled further to a final protein-level false discovery rate of 1%. Proteins were quantified by summing reporter ion counts across all matching PSMs, also as described previously^44^. Reporter ion intensities were adjusted to correct for the isotopic impurities of the different TMTpro reagents according to manufacturer specifications. The signal-to-noise measurements of peptides assigned to each protein were summed and these values were normalized so that the sum of the signal for all proteins in each channel was equivalent to account for equal protein loading. Finally, each protein abundance measurement was scaled, such that the summed signal-to-noise for that protein across all channels equals 100, thereby generating a relative abundance measurement.

MSstatsTMT was performed on peptides with >200 summed signal-to-noise ratios across TMT channels. For each protein, the filtered PSM TMTpro raw intensities were summed and log_2_ normalized to calculate protein quantification values (weighted average) and normalized to total TMT channel intensity across all quantified PSMs (adjusted to median total TMT intensity for the TMT channels). The log_2_ normalized summed protein reporter intensities were compared using a Student’s *t*-test and *P* values were corrected for multiple hypotheses using the Benjamini–Hochberg adjustment.

---

**Figure S1.**
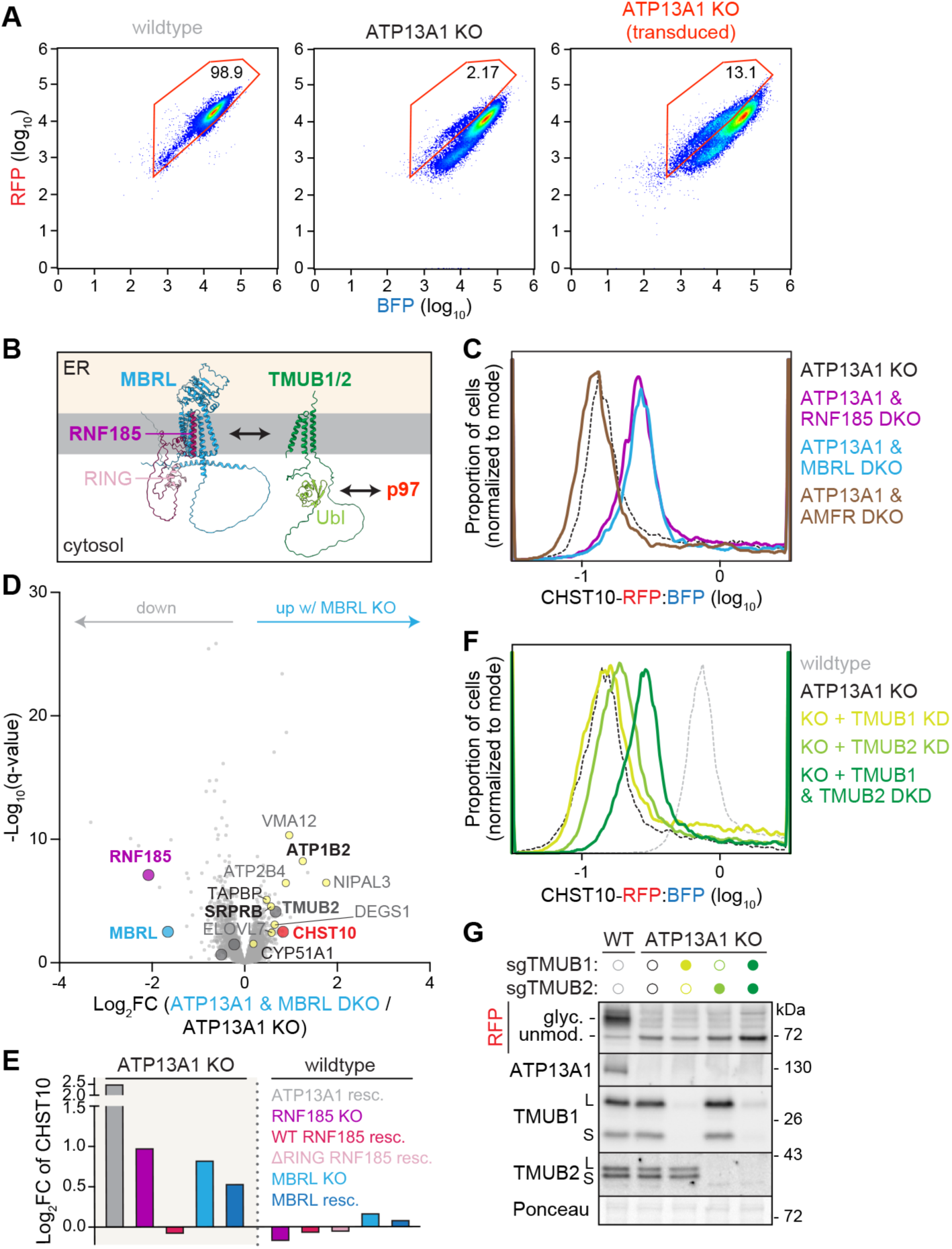
Factors that destabilize misoriented CHST10. **(A)** Fluorescent flow cytometry scatter plots of RFP vs. BFP levels in wildtype or ATP13A1 knockout (KO) Flp-In 293 T-REx cells expressing the CHST10 stability reporter without or with transduction (right) of the TKOv3 library. Percentage of cells within the gate is indicated. Note: TKOv3 library transduction increases the proportion of ATP13A1 KO cells with higher RFP:BFP levels. **(B)** Alphafold models of RNF185 (purple) in complex with MBRL (blue) and of TMUB1 (green). The RNF185 RING domain (light purple) and TMUB1 Ubl domain (light green) are indicated. **(C)** Knocking out RNF185 or MBRL stabilizes CHST10 in ATP13A1 KO cells. Fluorescent flow cytometry of the CHST10 stability reporter in wildtype (gray) or ATP13A1 KO (black) cells without or with additional knockout (double knockout, DKO) of RNF185 (purple), MBRL (light blue), or AMFR (brown). **(D)** TMT-MS volcano plot showing changes in protein levels upon knocking out MBRL (DKO) in ATP13A1 knockout (KO) Flp-In T-REx cells. **(E)** Fold-change (Log_2_FC) in CHST10 levels relative to ATP13A1 KO (left, shaded) or wildtype (right) cells upon ATP13A1 re-expression (gray), RNF185 KO without (purple) or with re-expression of WT (hot pink) or ΔRING (lavender) RNF185, or MBRL KO without (light blue) or with re-expression (dark blue) of MBRL. **(F)** Fluorescent flow cytometry of wildtype (gray) or ATP13A1 KO (black) cells expressing the CHST10 stability reporter without or with polyclonal sgRNA-mediated knockdown (KD) of TMUB1, TMUB2, or both (double knockdown, DKD). Note: knocking down both TMUB1 and TMUB2 stabilizes CHST10 in ATP13A1 KO cells. **(G)** SDS-PAGE and immunoblotting of lysates from cells as in (F).

**Figure S2.**
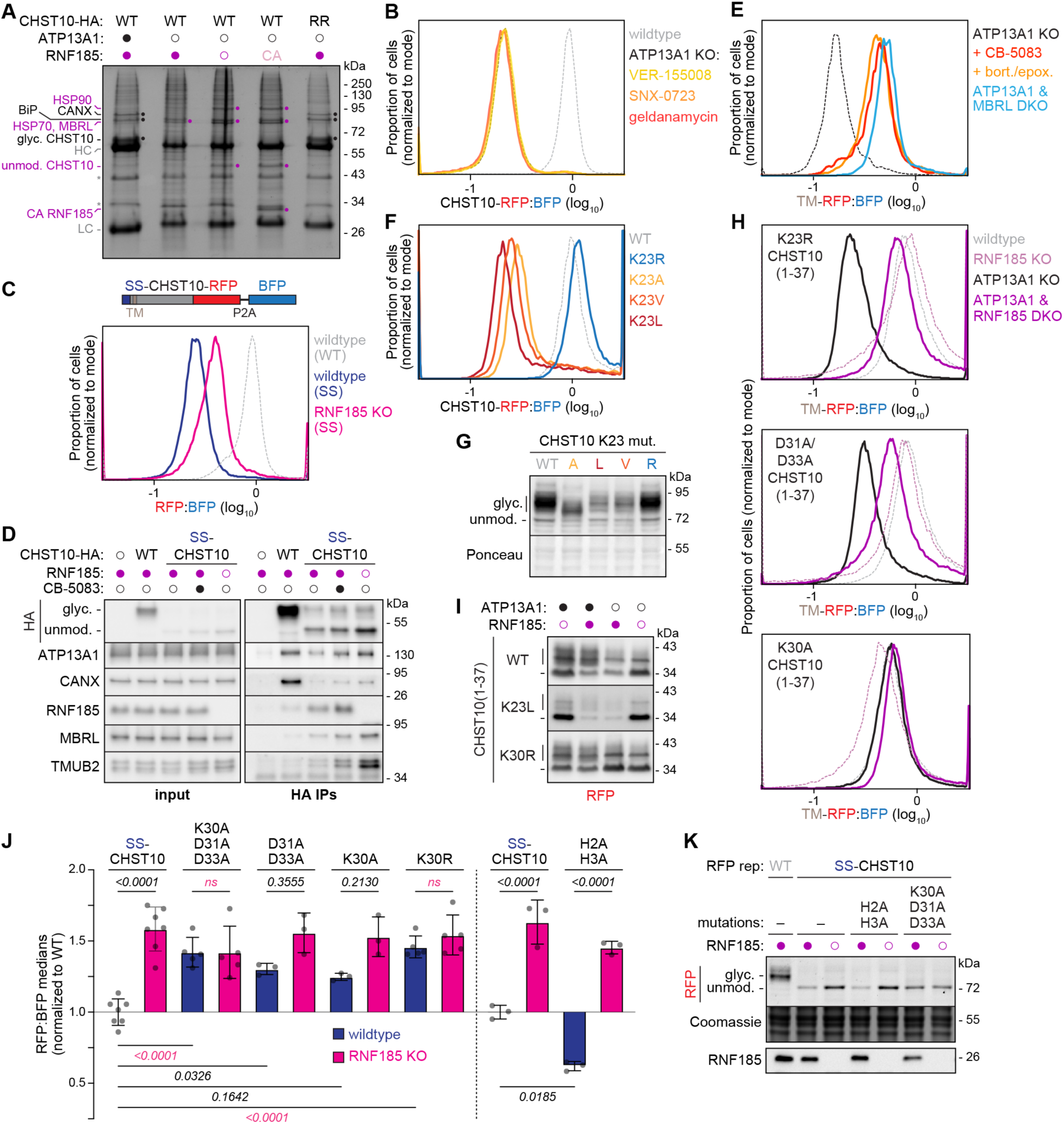
TM misorientation drives CHST10 destabilization. **(A)** Misoriented CHST10 specifically interacts with the RNF185 complex. SDS-PAGE and Coomassie staining of affinity purifications of wildtype (WT) or H2R/H3R (RR) C-terminally HA-tagged CHST10 in cells without (filled circles) or with knockout of ATP13A1 (open black circles) and/or RNF185 (open purple circles), and with re-expression of catalytically inactive C76A/C79A (CA) RNF185 (lane 4). Glyc. and unmod. CHST10 and near-stoichiometric proteins, identified by mass spectrometry, that co-elute with correctly oriented (black dots and labels) or misoriented (purple dots and labels) CHST10, are indicated. HC – heavy chain, LC – light chain, * – nonspecific interactors of antibody. **(B)** Fluorescent flow cytometry histograms of the RFP:BFP ratios of the CHST10 stability reporter in wildtype (left) or ATP13A1 KO (right) cells without or with treatment for 7 hr with inhibitors against HSP70 (30 µM VER155008) or HSP90 (500 nM SNX-0723, 2 µM geldanamycin). **(C)** Scheme (top) of the SS-CHST10-RFP stability reporter with the N-terminal cleavable SS of preprolactin appended to CHST10 and fluorescent flow cytometry (bottom) of wildtype or RNF185 KO Flp-In 293 T-REx cells expressing the wildtype (WT) CHST10 or the SS-CHST10 stability reporter. **(D)** SDS-PAGE and immunoblotting of lysates before (bottom) or after anti-HA immunoprecipitations (IPs, top) of C-terminally HA-tagged WT CHST10 or SS-CHST10 in wildtype (filled purple circles) or RNF185 KO (open purple circles) cells treated without or with 1 µM CB-5083 (filled black circle) for 6 hr. Note: SS-CHST10 is not glycosylated and specifically interacts with the RNF185 complex. **(E)** Fluorescent flow cytometry of ATP13A1 knockout (KO) without or with additional MBRL KO (blue) Flp-In 293 T-REx cells expressing a stability reporter containing the first 37 amino acids of CHST10 appended to RFP (TM-RFP) and separated from BFP by a P2A ribosome skipping sequence, treated for 6 hr without (black) or with p97 AAA-ATPase (1 µM CB-5083, dark orange) or proteasome (0.5 µM bortezomib, bort., and 0.5 µM epoxomicin, epox., light orange) inhibitors. Note: the CHST10 TM reporter in ATP13A1 KO cells is stabilized by the inhibitors and by MBRL KO. **(F)** Fluorescent flow cytometry histograms of the RFP:BFP ratios of the full-length CHST10 stability reporter in wildtype cells without (WT, gray) or with mutation of K23 to arginine (blue), alanine (light orange), valine (orange), or leucine (dark red). **(G)** SDS-PAGE and immunoblotting of lysates from cells as in (F). Note: K23A, K23L, and K23V all show reduced levels of glycosylated CHST10. Because K23A displays a different glycosylation pattern, we focused analysis on K23L. **(H)** Fluorescent flow cytometry of the RFP:BFP ratios of the minimal CHST10-TM stability reporter with the indicated mutation(s) in the indicated cell lines. **(I)** SDS-PAGE and immunoblotting for RFP of lysates from cells without (filled circles) or with (open circles) ATP13A1 and/or RNF185 KO expressing the indicated CHST10-TM reporters. Note: the K23L mutation reduces reporter glycosylation (vertical line) and independently destabilizes the reporter in wildtype cells, while the K30R mutation stabilizes unmodified reporter (dash) populations in all cell lines. **(J)** Individual (dots) and average (bars) median RFP:BFP ratios +/- s.d. from >3 independent fluorescent flow cytometry measurements of the indicated SS-CHST10 stability reporter variants in wildtype (navy) or RNF185 KO (pink) cells. P-values (one-way ANOVA) between the values in wildtype and RNF185 KO cells for each variant (top) and between SS-CHST10 and each variant in wildtype cells (bottom) are indicated. Measurements of the H2A/H3A variant were collected with different laser settings and are compared to measurements of the SS-CHST10 reporter collected under the same conditions. **(K)** SDS-PAGE and in-gel RFP fluorescence and Coomassie staining (top) or immunoblotting (bottom) of lysates from wildtype (filled circles) or RNF185 KO (open circles) cells expressing WT CHST10 or the indicated SS-CHST10 variant reporter.

**Figure S3.**
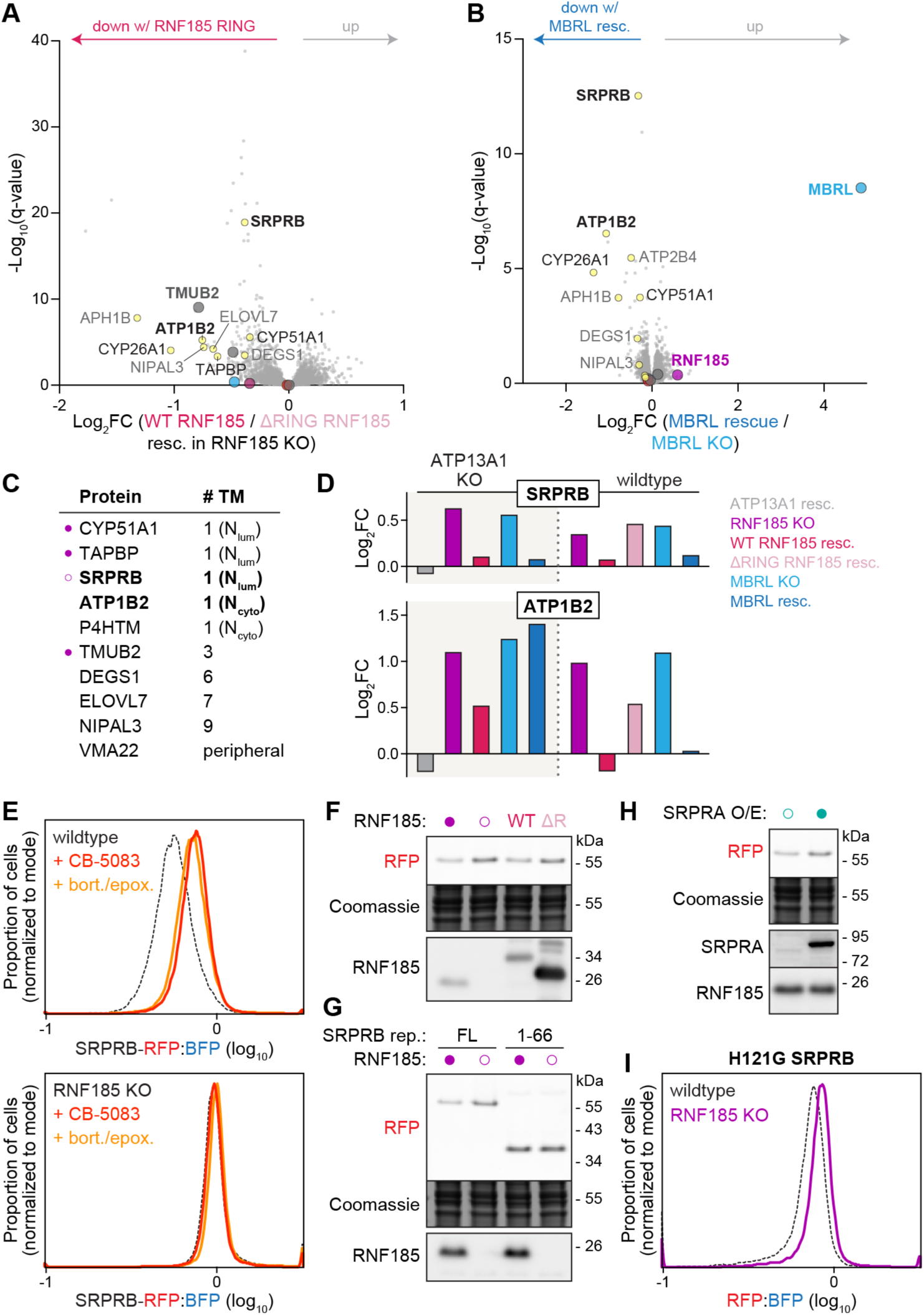
Identification of membrane proteins destabilized by the RNF185 complex. **(A)** TMT-MS volcano plot showing differences in protein levels in RNF185 KO cells re-expressing either wildtype (WT) or ΔRING Flag-tagged RNF185. Colored dots indicate RNF185 (purple), MBRL (light blue), CHST10 (red), and other RNF185-sensitive proteins (yellow). **(B)** TMT-MS volcano plot showing changes in protein levels upon re-expressing Flag-tagged MBRL (rescue) in MBRL KO Flp-In 293 T-REx cells. **(C)** Endogenous proteins identified to be destabilized by RNF185 in both wildtype and ATP13A1 KO backgrounds, with their predicted number of transmembrane helices (# TM) and topology for single-spanning proteins. Previously noted RNF185 substrates (filled purple circles) and interactors (open circle) are indicated. **(D)** Fold-change (Log_2_FC) in the levels of SRPRB (top) or ATP1B2 (bottom) relative to ATP13A1 KO (left) or wildtype (right) cells upon ATP13A1 re-expression (resc., gray), RNF185 KO without (purple) or with re-expression of WT (hot pink) or ΔRING (lavender) RNF185, or MBRL KO without (light blue) or with re-expression (dark blue) of MBRL. **(E)** Fluorescent flow cytometry histograms of the RFP:BFP ratios of a stability reporter of RFP-tagged SRPRB, separated from BFP by a P2A ribosome skipping sequence, in wildtype (top) or RNF185 knockout (KO, bottom) Flp-In 293 T-REx cells without (black) or with inhibitors of p97 AAA-ATPase (1 µM CB-5083, dark orange) or proteasome (0.5 µM bortezomib, bort., and 0.5 µM epoxomicin, epox., light orange) activity for 6 hr. **(F)** SDS-PAGE and in-gel RFP fluorescence and Coomassie staining (top) or immunoblotting (bottom) of the SRPRB stability reporter in wildtype (filled circle) or RNF185 KO cells without (open circle) or with re-expression of WT or ΔRING (ΔR) Flag-tagged RNF185. **(G)** As in (F) of stability reporters of full-length (FL) or the first 66 amino acids (1-66) of SRPRB in wildtype (filled circles) or RNF185 KO (open circles) cells. **(H)** As in (F) of the SRPRB stability reporter in wildtype cells without (open circle) or with (filled circle) SRPRA overexpression (O/E). **(I)** Fluorescent flow cytometry of the H121G SRPRB stability reporter in wildtype (black) or RNF185 KO (purple) cells.

**Figure S4.**
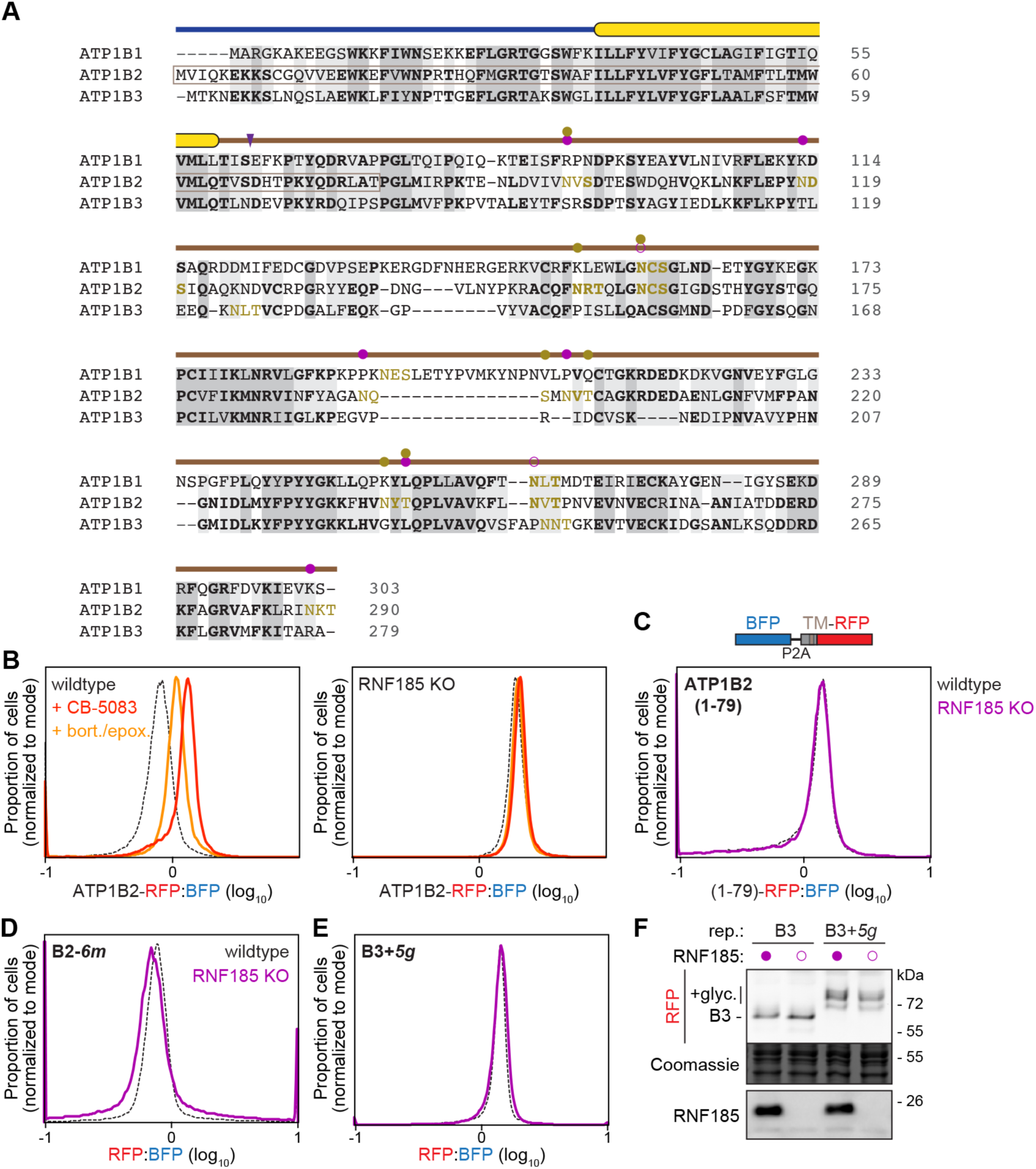
Analysis of ATP1B paralog variants. **(A)** Sequence alignment of ATP1B1, ATP1B2, and ATP1B3, with the cytosolic N-terminal domain (dark blue line), predicted transmembrane helix (yellow), and lumenal domain (brown) indicated above. The first 79 amino acids of ATP1B2 (boxed) used for the TM reporter, the chimera switch site (arrowhead), and N-linked glycosylation motifs (gold) are shown. Purple and gold dots denote the locations of loss-of and gain-of glycosylation mutations made in ATP1B2 and ATP1B3, respectively. Open purple dots – mutations only in the ATP1B2-*8m* mutant. **(B)** Fluorescent flow cytometry histograms of the RFP:BFP ratios of a stability reporter of RFP-tagged ATP1B2 following BFP and a P2A ribosome skipping sequence in wildtype (left) or RNF185 KO (right) cells without (black) or with inhibitors of p97 AAA-ATPase (1 µM CB-5083, dark orange) or proteasome (0.5 µM bortezomib, bort., and 0.5 µM epoxomicin, epox., light orange) activity for 6 hr. **(C)** Scheme (top) and fluorescent flow cytometry histograms of the RFP:BFP ratios (bottom) of a stability reporter of the first 79 amino acids of ATP1B2 (1-79) in wildtype (black) or RNF185 KO (purple) cells. Note: this reporter is not sensitive to RNF185. **(D)** Fluorescent flow cytometry histograms of the RFP:BFP ratios of the ATP1B2-*6m* stability reporter in which six N-linked glycosylation motifs in ATP1B2 are mutated. **(E)** Fluorescent flow cytometry of the RFP:BFP ratios of the ATP1B3+*5g* stability reporter in which five N-linked glycosylation motifs are added to ATP1B3. **(F)** SDS-PAGE and in-gel RFP fluorescence and Coomassie staining (top) or immunoblotting (bottom) of lysates from wildtype (filled circles) or RNF185 KO (open circles) cells expressing the indicated RFP-tagged stability reporters of ATP1B3 without or with (ATP1B3+*5g*) mutations to add five N-linked glycosylation sites.

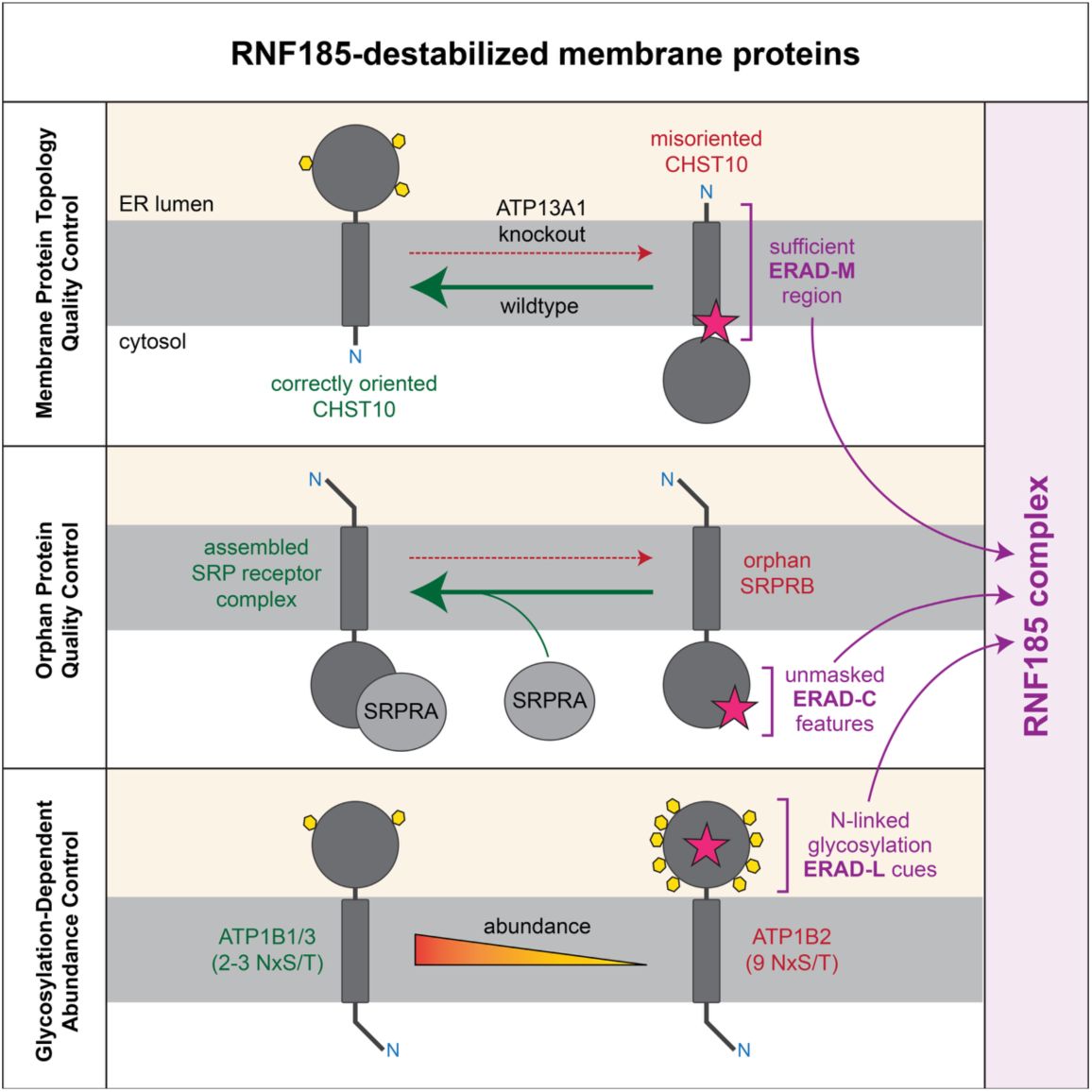

## Notes

### Competing Interest Statement

The authors have declared no competing interest.

## References

1. Christianson, J.C., Jarosch, E., and Sommer, T. (2023). Mechanisms of substrate processing during ER-associated protein degradation. Nat. Rev. Mol. Cell Biol. 24, 777–796. 10.1038/s41580-023-00633-8.

2. Olzmann, J.A., Kopito, R.R., and Christianson, J.C. (2013). The mammalian endoplasmic reticulum-associated degradation system. Cold Spring Harb. Perspect. Biol. 5, a013185– a013185. 10.1101/cshperspect.a013185.

3. Wu, X., Siggel, M., Ovchinnikov, S., Mi, W., Svetlov, V., Nudler, E., Liao, M., Hummer, G., and Rapoport, T.A. (2020). Structural basis of ER-associated protein degradation mediated by the Hrd1 ubiquitin ligase complex. Science 368, eaaz2449. 10.1126/science.aaz2449.

4. Schmidt, C.C., Vasic, V., and Stein, A. (2020). Doa10 is a membrane protein retrotranslocase in ER-associated protein degradation. Elife 9, e56945. 10.7554/elife.56945.

5. Baldridge, R.D., and Rapoport, T.A. (2016). Autoubiquitination of the Hrd1 Ligase Triggers Protein Retrotranslocation in ERAD. Cell 166, 394–407. 10.1016/j.cell.2016.05.048.

6. Carvalho, P., Goder, V., and Rapoport, T.A. (2006). Distinct Ubiquitin-Ligase Complexes Define Convergent Pathways for the Degradation of ER Proteins. Cell 126, 361–373.

7. Huyer, G., Piluek, W.F., Fansler, Z., Kreft, S.G., Hochstrasser, M., Brodsky, J.L., and Michaelis, S. (2004). Distinct Machinery Is Required in Saccharomyces cerevisiae for the Endoplasmic Reticulum-associated Degradation of a Multispanning Membrane Protein and a Soluble Luminal Protein. J. Biol. Chem. 279, 38369–38378. 10.1074/jbc.m402468200.

8. Vashist, S., and Ng, D.T.W. (2004). Misfolded proteins are sorted by a sequential checkpoint mechanism of ER quality control. J. Cell Biol. 165, 41–52. 10.1083/jcb.200309132.

9. Sato, B.K., Schulz, D., Do, P.H., and Hampton, R.Y. (2009). Misfolded membrane proteins are specifically recognized by the transmembrane domain of the Hrd1p ubiquitin ligase. Mol. Cell 34, 212–222. 10.1016/j.molcel.2009.03.010.

10. Russ, J.E., Peterson, B.G., Taylor, S., and Baldridge, R.D. (2025). Competition between Der1 and ERAD-M substrates controls Hrd1 complex function. Proc. Natl. Acad. Sci. 122, e2513595122. 10.1073/pnas.2513595122.

11. Bordallo, J., Plemper, R.K., Finger, A., and Wolf, D.H. (1998). Der3p/Hrd1p Is Required for Endoplasmic Reticulum-associated Degradation of Misfolded Lumenal and Integral Membrane Proteins. Mol. Biol. Cell 9, 209–222. 10.1091/mbc.9.1.209.

12. Habeck, G., Ebner, F.A., Shimada-Kreft, H., and Kreft, S.G. (2015). The yeast ERAD-C ubiquitin ligase Doa10 recognizes an intramembrane degron. J. Cell Biol. 209, 261–273. 10.1083/jcb.201408088.

13. McKenna, M.J., Adams, B.M., Chu, V., Paulo, J.A., and Shao, S. (2022). ATP13A1 prevents ERAD of folding-competent mislocalized and misoriented proteins. Mol. Cell 82, 4277–4289.e10. 10.1016/j.molcel.2022.09.035.

14. Stefanovic-Barrett, S., Dickson, A.S., Burr, S.P., Williamson, J.C., Lobb, I.T., Boomen, D.J., Lehner, P.J., and Nathan, J.A. (2018). MARCH6 and TRC8 facilitate the quality control of cytosolic and tail-anchored proteins. EMBO Rep. 19. 10.15252/embr.201745603.

15. Christianson, J.C., Olzmann, J.A., Shaler, T.A., Sowa, M.E., Bennett, E.J., Richter, C.M., Tyler, R.E., Greenblatt, E.J., Harper, J.W., and Kopito, R.R. (2012). Defining human ERAD networks through an integrative mapping strategy. Nat. Cell Biol. 14, 93–105. 10.1038/ncb2383.

16. Fenech, E.J., Lari, F., Charles, P.D., Fischer, R., Laétitia-Thézénas, M., Bagola, K., Paton, A.W., Paton, J.C., Gyrd-Hansen, M., Kessler, B.M., et al. (2020). Interaction mapping of endoplasmic reticulum ubiquitin ligases identifies modulators of innate immune signalling. Elife 9, e57306. 10.7554/elife.57306.

17. Weijer, M.L. van de, Krshnan, L., Liberatori, S., Guerrero, E.N., Robson-Tull, J., Hahn, L., Lebbink, R.J., Wiertz, E.J.H.J., Fischer, R., Ebner, D., et al. (2020). Quality Control of ER Membrane Proteins by the RNF185/Membralin Ubiquitin Ligase Complex. Mol. Cell 79, 768–781.e7. 10.1016/j.molcel.2020.07.009.

18. Weijer, M.L. van de, Samanta, K., Sergejevs, N., Jiang, L., Dueñas, M.E., Heunis, T., Huang, T.Y., Kaufman, R.J., Trost, M., Sanyal, S., et al. (2024). Tapasin assembly surveillance by the RNF185/Membralin ubiquitin ligase complex regulates MHC-I surface expression. Nat. Commun. 15, 8508. 10.1038/s41467-024-52772-x.

19. Sergejevs, N., Avci, D., Weijer, M.L. van de, Corey, R.A., Lemberg, M.K., and Carvalho, P. (2024). Topology surveillance of the lanosterol demethylase CYP51A1 by signal peptide peptidase. J. Cell Sci. 137, jcs262333. 10.1242/jcs.262333.

20. Zhu, B., Jiang, L., Huang, T., Zhao, Y., Liu, T., Zhong, Y., Li, X., Campos, A., Pomeroy, K., Masliah, E., et al. (2017). ER-associated degradation regulates Alzheimer’s amyloid pathology and memory function by modulating γ-secretase activity. Nat. Commun. 8, 1472. 10.1038/s41467-017-01799-4.

21. Heijne, G. von (1989). Control of topology and mode of assembly of a polytopic membrane protein by positively charged residues. Nature 341, 456–458. 10.1038/341456a0.

22. Wahlberg, J.M., and Spiess, M. (1997). Multiple Determinants Direct the Orientation of Signal–Anchor Proteins: The Topogenic Role of the Hydrophobic Signal Domain. J. Cell Biol. 137, 555–562. 10.1083/jcb.137.3.555.

23. McKenna, M.J., Sim, S.I., Ordureau, A., Wei, L., Harper, J.W., Shao, S., and Park, E. (2020). The endoplasmic reticulum P5A-ATPase is a transmembrane helix dislocase. Science 369. 10.1126/science.abc5809.

24. Wang, L., Li, J., Wang, Q., Ge, M.-X., Ji, J., Liu, D., Wang, Z., Cao, Y., Zhang, Y., and Zhang, Z.-R. (2022). TMUB1 is an endoplasmic reticulum-resident escortase that promotes the p97-mediated extraction of membrane proteins for degradation. Mol. Cell 82, 3453–3467.e14. 10.1016/j.molcel.2022.07.006.

25. Rapoport, T.A., Li, L., and Park, E. (2017). Structural and Mechanistic Insights into Protein Translocation. Annu. Rev. Cell Dev. Biol. 33, 369–390. 10.1146/annurev-cellbio-100616-060439.

26. Schwartz, T., and Blobel, G. (2003). Structural Basis for the Function of the β Subunit of the Eukaryotic Signal Recognition Particle Receptor. Cell 112, 793–803. 10.1016/s0092-8674(03)00161-2.

27. Ogg, S.C., Barz, W.P., and Walter, P. (1998). A Functional GTPase Domain, but not its Transmembrane Domain, is Required for Function of the SRP Receptor β-subunit. J. Cell Biol. 142, 341–354. 10.1083/jcb.142.2.341.

28. Zou, C., Yoon, H., Park, P.M.C., Patten, J.J., Pellman, J., Carreiro, J., Tsai, J.M., Li, Y.-D., Burman, S.S.R., Donovan, K.A., et al. (2023). The human E3 ligase RNF185 is a regulator of the SARS-CoV-2 envelope protein. iScience 26, 106601. 10.1016/j.isci.2023.106601.

29. Cho, N.H., Cheveralls, K.C., Brunner, A.-D., Kim, K., Michaelis, A.C., Raghavan, P., Kobayashi, H., Savy, L., Li, J.Y., Canaj, H., et al. (2022). OpenCell: Endogenous tagging for the cartography of human cellular organization. Science 375, eabi6983. 10.1126/science.abi6983.

30. Itzhak, D.N., Tyanova, S., Cox, J., and Borner, G.H. (2016). Global, quantitative and dynamic mapping of protein subcellular localization. Elife 5, e16950. 10.7554/elife.16950.

31. Ravid, T., Kreft, S.G., and Hochstrasser, M. (2006). Membrane and soluble substrates of the Doa10 ubiquitin ligase are degraded by distinct pathways. EMBO J. 25, 533–543. 10.1038/sj.emboj.7600946.

32. Bernasconi, R., Galli, C., Calanca, V., Nakajima, T., and Molinari, M. (2010). Stringent requirement for HRD1, SEL1L, and OS-9/XTP3-B for disposal of ERAD-LS substrates. J. Cell Biol. 188, 223–235. 10.1083/jcb.200910042.

33. Khouri, E.E., Pavec, G.L., Toledano, M.B., and Delaunay-Moisan, A. (2013). RNF185 Is a Novel E3 Ligase of Endoplasmic Reticulum-associated Degradation (ERAD) That Targets Cystic Fibrosis Transmembrane Conductance Regulator (CFTR). J. Biol. Chem. 288, 31177– 31191. 10.1074/jbc.m113.470500.

34. Riepe, C., Wąchalska, M., Deol, K.K., Amaya, A.K., Porteus, M.H., Olzmann, J.A., and Kopito, R.R. (2024). Small-molecule correctors divert CFTR-F508del from ERAD by stabilizing sequential folding states. Mol. Biol. Cell 35, ar15. 10.1091/mbc.e23-08-0336.

35. Qiu, D., Wang, Q., Wang, Z., Chen, J., Yan, D., Zhou, Y., Li, A., Zhang, R., Wang, S., and Zhou, J. (2018). RNF185 modulates JWA ubiquitination and promotes gastric cancer metastasis. Biochim. Biophys. Acta Mol. Basis Dis. 1864, 1552–1561. 10.1016/j.bbadis.2018.02.013.

36. Zhou, Y., Shang, H., Zhang, C., Liu, Y., Zhao, Y., Shuang, F., Zhong, H., Tang, J., and Hou, S. (2014). The E3 ligase RNF185 negatively regulates osteogenic differentiation by targeting Dvl2 for degradation. Biochem. Biophys. Res. Commun. 447, 431–436. 10.1016/j.bbrc.2014.04.005.

37. Zhang, J., Wang, B., Gao, X., Peng, C., Shan, C., Johnson, S.F., Schwartz, R.C., and Zheng, Y.-H. (2022). RNF185 regulates proteostasis in Ebolavirus infection by crosstalk between the calnexin cycle, ERAD, and reticulophagy. Nat. Commun. 13, 6007. 10.1038/s41467-022-33805-9.

38. Wang, Q., Huang, L., Hong, Z., Lv, Z., Mao, Z., Tang, Y., Kong, X., Li, S., Cui, Y., Liu, H., et al. (2017). The E3 ubiquitin ligase RNF185 facilitates the cGAS-mediated innate immune response. PLoS Pathog. 13, e1006264. 10.1371/journal.ppat.1006264.

39. Zhang, J., Lu, X., Li, S., Wang, T., Ahmad, I., and Zheng, Y. (2026). Membralin Assembles a MAN1B1–VCP Complex to Target Foreign Glycoproteins from the Endoplasmic Reticulum to Lysosomes for Degradation. Adv. Sci. 13, e19256. 10.1002/advs.202519256.

40. Yang, B., Qu, M., Wang, R., Chatterton, J.E., Liu, X.-B., Zhu, B., Narisawa, S., Millan, J.L., Nakanishi, N., Swoboda, K., et al. (2015). The critical role of membralin in postnatal motor neuron survival and disease. Elife 4, e06500. 10.7554/elife.06500.

41. Hart, T., Tong, A.H.Y., Chan, K., Leeuwen, J.V., Seetharaman, A., Aregger, M., Chandrashekhar, M., Hustedt, N., Seth, S., Noonan, A., et al. (2017). Evaluation and Design of Genome-Wide CRISPR/SpCas9 Knockout Screens. G3 *7*, 2719–2727. 10.1534/g3.117.041277.

42. Li, W., Xu, H., Xiao, T., Cong, L., Love, M.I., Zhang, F., Irizarry, R.A., Liu, J.S., Brown, M., and Liu, X.S. (2014). MAGeCK enables robust identification of essential genes from genome-scale CRISPR/Cas9 knockout screens. Genome Biol. 15, 554. 10.1186/s13059-014-0554-4.

43. Huttlin, E.L., Jedrychowski, M.P., Elias, J.E., Goswami, T., Rad, R., Beausoleil, S.A., Villén, J., Haas, W., Sowa, M.E., and Gygi, S.P. (2010). A tissue-specific atlas of mouse protein phosphorylation and expression. Cell 143, 1174–1189. 10.1016/j.cell.2010.12.001.

44. McAlister, G.C., Huttlin, E.L., Haas, W., Ting, L., Jedrychowski, M.P., Rogers, J.C., Kuhn, K., Pike, I., Grothe, R.A., Blethrow, J.D., et al. (2012). Increasing the Multiplexing Capacity of TMTs Using Reporter Ion Isotopologues with Isobaric Masses. Anal. Chem. 84, 7469–7478. 10.1021/ac301572t.

